# Cellular behavior analysis from live-cell imaging of TCR T cell–cancer cell interactions

**DOI:** 10.1101/2024.11.19.624390

**Authors:** Archit Verma, Changhua Yu, Stefanie Bachl, Ivan Lopez, Morgan Schwartz, Erick Moen, Nupura Kale, Carter Ching, Geneva Miller, Tom Dougherty, Ed Pao, William Graf, Carl Ward, Siddhartha Jena, Alex Marson, Julia Carnevale, David Van Valen, Barbara E Engelhardt

## Abstract

T cell therapies, such as chimeric antigen receptor (CAR) T cells and T cell receptor (TCR) T cells, are a growing class of anti-cancer treatments. However, expansion to novel indications and beyond last-line treatment requires engineering cells’ dynamic population behaviors. Here we develop the tools for *cellular behavior analysis* of T cells from live-cell imaging, a common and inexpensive experimental setup used to evaluate engineered T cells. We first develop a state-of-the-art segmentation and tracking pipeline, *Caliban*, based on human-in-the-loop deep learning. We then build the *Occident* pipeline to collect a catalog of phenotypes that characterize cell populations, morphology, movement, and interactions in co-cultures of modified T cells and antigen-presenting tumor cells. We use Caliban and Occident to interrogate how interactions between T cells and cancer cells differ when beneficial knock-outs of *RASA2* and *CUL5* are introduced into TCR T cells. We apply spatiotemporal models to quantify T cell recruitment and proliferation after interactions with cancer cells. We discover that, compared to a safe harbor knockout control, *RASA2* knockout T cells have longer interaction times with cancer cells leading to greater T cell activation and killing efficacy, while *CUL5* knockout T cells have increased proliferation rates leading to greater numbers of T cells for hunting. Together, segmentation and tracking from Caliban and phenotype quantification from Occident enable cellular behavior analysis to better engineer T cell therapies for improved cancer treatment.

## Introduction

Chimeric antigen receptor (CAR) T cells are a groundbreaking therapy for haemotological cancers, and T cell receptor (TCR)-transgenic T cells are beginning to be used to treat solid tumors^1,2^. These therapies currently use *ex vivo* transduction and expansion of a patient’s T cells to express surface proteins that recognize cancerous cells and activate T cells, boosting the immune system’s anti-cancer response^3^. CAR T cells are now standard treatment for a subset of B cell leukemias and lymphomas, particularly those that return after traditional chemotherapy, where few treatment options previously existed; TCR T cell treatments are now available for metastatic synovial sarcoma^2^. Clinicians are optimistic about moving these engineered T cell therapies up from a last-line option to an alternative for chemotherapy or radiation. The antigen-specificity of CAR T and related treatments should have fewer off-target effects than traditional cancer therapies and allow tuning to surface markers that target a wide variety of cancer cells with high precision. However, challenges remain in engineering both novel T cell therapies for new indications and making current T cell therapies safer for general use. For example, CAR T therapy today can have severe side effects such as cytokine release syndrome (CRS)^4,5^, a dysregulated immune response that can cause marked toxicity. Expanded use to solid tumors will also require overcoming the complex spatial tumor microenvironment that suppresses immune activity physically and chemically^1,6^.

Genetic editing is a promising approach to engineering novel immune cancer therapies that may overcome these challenges^7–11^. The recent development of a suite of CRISPR-Cas based editing tools allows tunable knockouts, knockins, inhibitions, targeted mutations, and epigenetic alterations of the genes in a T cell, providing unprecedented opportunities to control T cell functions in a tunable manner. High-throughput screening protocols have uncovered genetic modifications that show beneficial effects on T cell anti-cancer behavior in terms of proliferation, molecular signaling, and mouse model efficacy^7,12–14^. For example, ablation of *RASA2*, a gene that codes for a signaling checkpoint in human T cells, sensitizes T cells to antigen-stimulation, even in the presence of various immunosuppressive challenges, and makes T cells more resilient to repetitive antigen stimulation^13^. Ablation of *CUL5* in T cells, another negative regulator of T cell signaling via the Cullin-5 complex, also improved immune inhibition of cancer growth in prior studies^7,8^. These knockouts are two of the many candidate genetic perturbations identified by single-phenotype screens, including cancer cell proliferation or cytokine assays. However, the translation of genetic perturbations identified by single-phenotype screens to effective *in vivo* cancer cell killing in mice and humans is often poor. This may be due to the narrow readouts most assays provide, focusing on single measurements that are not representative of the complex spatiotemporal group dynamics and behaviors of T cells and T cell interactions with cancer cells.

Engineering effective cell therapies that translate from bench to bedside will require *cellular behavioral analyses* to characterize the complex dynamics of *in vivo* systems. Similar to studies of the response of individual mice^15–17^ or populations of animals^18–20^ to various stimuli, interrogating the behavioral responses of individual and populations of cells under conditions such as genetic modifications unlocks a deeper understanding of the multiscale biology at play. Comprehensive cellular behavioral analyses will require data collection on populations of cells across space and time, and the development of associated computational and statistical techniques to interrogate such data. The tools and techniques to begin such work, however, are already available.

Unlike the static, single time-point readouts from single-cell RNA-seq or FISH-based spatial readouts, *live-cell imaging* holds the potential to capture these complex behaviors across time and space^21^. Brightfield and fluorescent imaging is already a ubiquitous method to capture T cell efficacy through single-phenotype analyses. But these data also hold the potential for measuring more complex behavioral phenotypes. Live-cell imaging of fluorescence-labeled T cells profiled serial killing^22–24^ and cellular cooperation^25^ during treatment, among other complex phenotypes^21^. Novel three-dimensional imaging systems have served as the basis for engineering T cell response to cancerous organoids^24^ or modulating T cell velocity to infiltrate solid tumors^26^. Nevertheless, three-dimensional imaging is expensive and inaccessible for many studies, while fluorescently-labeled T cell lines are difficult to create, hard to generalize across T cell subpopulations, and can alter T cell behavior, limiting accurate and scalable behavioral phenotyping.

Imaging a two-dimensional plane with only fluorescent-tagged cancer nuclei is a cheaper alternative that is commonly used in T cell engineering^12,13^ (Figure 1a). T cells with a variety of modifications can be easily and scalably co-cultured with antigen-expressing cancer cells to understand how each modification affects cancer cell killing. The total fluorescence from a cancer nuclear marker over time is the single phenotype used to summarize the anti-cancer effect of a modified T cell. However, an enormous amount of biologically important and actionable information about the modified T cells and their interaction with cancer cells is lost in this simple analysis. Observation of brightfield videos shows a multitude of T cell and cancer cells behaviors such as interactions, aggregation, swarming, proliferation, cell death, and recruitment that goes beyond the total fluorescence coverage of cancer cells. Dynamics occur between and among cell populations, including but not limited to T cell activation as they bind to tumor cells, T cell recruitment of unbound T cells, and cancer cell aggregation as they die or try to evade the pack hunting behavior of the T cells. Cellular behavior analysis can capture dynamics such as population growth, cell movement, morphological changes upon interaction, and activation-induced recruitment and proliferation (Figure 1b). Engineering optimal and targeted T cell therapies will require an understanding and a direct quantification of these dynamics to overcome the multi-cellular nature of challenges such as mitigating cytokine release syndrome and successfully breaching tumor immune microenvironments.

**Figure 1.**
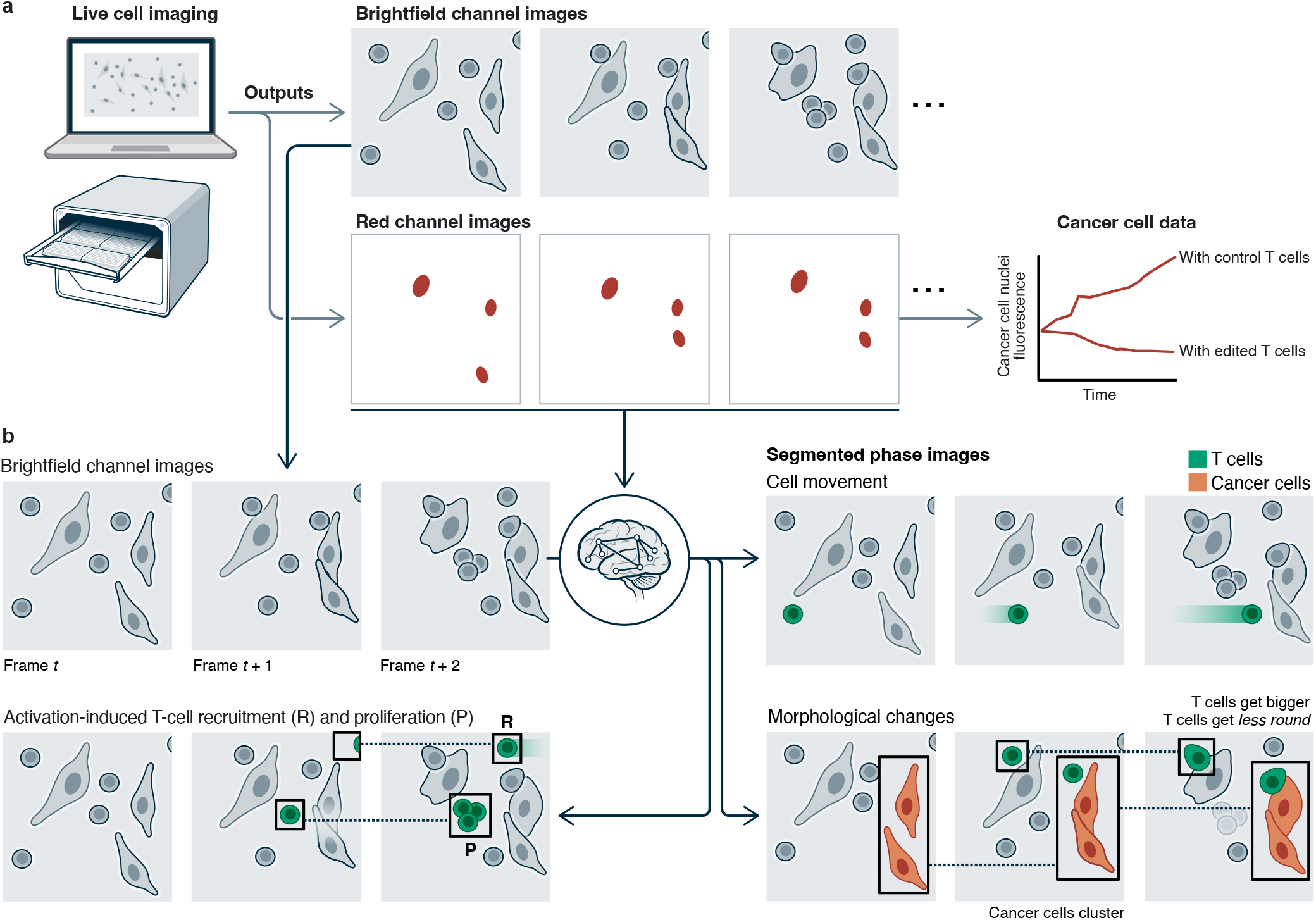
Live-cell imaging serves as a platform for cellular behavior analysis. (a) Traditional analysis of immunotherapy and tumor co-cultures monitors the fluorescence of a cancer nuclear marker over time to quantify killing rates, missing dynamic behaviors of T cells and tumor cells visible in brightfield images. (b) With novel machine learning methods for segmentation, tracking, and spatiotemporal modeling, brightfield images can be used to phenotype cellular behavior, particularly interaction dynamics between T cells and cancer cells.

Modern machine learning methods offer a compelling path to general solutions for cellular behavior analyses from imaging data. High-quality cellular tracking, identifying the same cell across different time points, is particularly critical to establishing the temporal aspects of cell behavior. Progress in deep learning solutions to cell tracking has been limited due to a lack of similar data resources and methodology for dynamic data. Existing datasets are limited in their scope and scale^27–34^, whereas simulated datasets have not yet proven capable of creating high-performing models^27,35,36^. Further, existing datasets are limited in the resolution of their labels (e.g., point labels versus pixel-level segmentation labels), trajectory length (the number of frames over which a cell is tracked), and the number of mitotic events. These limitations are understandable, given the time-consuming nature of labeling dynamic movies. Not only must each cell be segmented in a temporally consistent way, but lineage information must also be captured by tracking cells over time and labeling cell division events. Existing labeling methodology that has proven scalable for static images has yet to be extended at scale to these dynamic datasets^37^.

In this work, we develop and demonstrate the feasibility and benefits of cellular behavior analysis. We first develop the *Caliban* pipeline, which enables rapid and accurate segmentation, tracking, and lineage construction of nuclear live-cell imaging data with no manual parameter tuning. We combined a human-in-the-loop approach to image labeling^38^, adapted to dynamic imaging data, a novel deep learning algorithm for cell tracking, and new benchmarks for cell tracking to create a new labeled reference dataset, DynamicNuclearNet, for cell tracking. We then develop the *Occident* pipeline for cellular behavior analysis given cell segmentation and tracking. We apply Occident to co-cultures of TCR T cells and antigen-presenting cancer cells. We extract a catalogue of phenotypes from inexpensive 2D live-cell imaging using Sartorious Incucyte with a brightfield phase channel and red fluorescent channel marking cancer cell nuclei. We demonstrate the pipeline on live-cell imaging data, imaged every four minutes, of three TCR T cell lines - a population of safe harbor control knockout (SH KO) of *AAVS1*, a population with *RASA2* knockout (RASA2 KO), and a population with *CUL5* knockout (CUL5 KO) - co-cultured with CD19-expressing A375 RFP^+^ cancer cells. We identify time-dependent morphological and speed changes across perturbations that quantify T cell activity and function. We use cell tracking to identify interactions between cancer cells and T cells and report how morphology and motility changes after intercellular interactions. Finally, we use interpretable spatiotemporal models to disaggregate T cell recruitment and proliferation in response to antigen activation. This work serves as a major step towards cellular behavior analysis to deeply phenotype T cells and cancer cells, which can be scaled to larger genetic screens to engineer immunotherapies for improved therapeutic efficacy and broad deployment.

## Results

We first describe the development of Caliban as a state-of-the-art segmentation and tracking tool for cellular images. Trained on a new DynamicNuclearNet dataset, Caliban outperforms benchmark models across segmentation and tracking methods. We then describe the application of Occident to perform cellular behavior analysis. We apply Occident to live-cell imaging of co-cultures of cancer cells and TCR T cells with *RASA2* knockout (RASA2 KO), *CUL5* knockout (CUL5 KO), and a control safe harbor *AAVS1* knockout (SH KO). *RASA2* encodes the T cell signaling checkpoint RAS GTPase-activating protein, while *CUL5* is a cell growth and proliferation regulator in the Cullin-RING ligase 5 multi-protein complex involved in cytokine signaling^8^. Built on Caliban’s segmentation and tracking, Occident quantifies a catalog of phenotypes for both cancer and T cells that allow comparisons of the effects of different T cell genetic modifications. We use Occident to uncover cellular-level responses to T cell-cancer cell interactions. Moreover, Occident uses an interpretable spatial Markov model to quantify T cell proliferation and recruitment from activation upon interaction.

### Caliban effectively segments, tracks, and phenotypes live-cell imaging data

The first major challenge in establishing cellular behaviors from live-cell imaging is identifying single cells in each time frame – *segmentation* – and then establishing the same cell’s position over time – *tracking*. We therefore developed Caliban, an integrated solution to segmentation and tracking. Caliban uses a tracking-by-detection approach in which cells are first identified in each frame by a deep learning model; these detections are then used to reconstruct a lineage tree that connects cells across frames and through cell division events. For the reconstruction of lineage trees, we use a deep learning model that encodes temporal dependencies for multiple features of each object and predicts the probability of a parent–child relationship that exists due to a cell division event between any pair of cells across frames^42^.

Accurate cell detection and segmentation are essential to producing faithful lineage reconstructions. In addition to prior data and models^38,43^, we built DynamicNuclearNet, a segmented and tracked dataset of fluorescently-labeled cell nuclei spanning five different cell lines. This dataset contains 647,322 unique nuclear segmentations assembled into over 16,501 trajectories with over 2,621 division events. Each trajectory begins at the cell’s appearance in the field of view (FOV) or birth as a daughter cell, and ends when the cell disappears by permanently leaving the FOV, dying, or dividing. While generating pixel-level masks for each cell is expensive compared to other types of labels (e.g., centroids or bounding boxes), these masks facilitate downstream analysis steps, such as quantifying signaling reporters or nuclear morphology. The 2,621 division events in our dataset surpass all previous annotation efforts that use nuclear segmentation masks (Table 1), which allows us to incorporate cell division detection into our deep-learning-based cell-tracking method. We combined prior work on cell segmentation^38,43^ with DynamicNuclearNet and a comprehensive benchmarking framework to train an accurate deep learning model for nuclear segmentation as part of Caliban (see Methods for details).

In Caliban, raw images are passed through the nuclear segmentation model to produce cell masks. These masks are used to extract features for each cell, while the centroids are used to construct an adjacency matrix to identify cells in close proximity (*<* 64 pixels, 41.6 *µ*m). These features and the adjacency matrix are fed into a neighborhood encoder model, which uses a graph attention network^39,40^ to generate feature vectors that summarize information about a cell’s—and its neighbors’—appearance, location, and morphology (Figure 2b). These feature vectors are then fed into a tracking model that causally integrates temporal information and performs a pairwise comparison of each cell’s feature vector across frames to produce an effective probability score indicating whether two cells are the same cell, are different cells, or have a parent–child relationship (Figure 2c).

**Figure 2.**
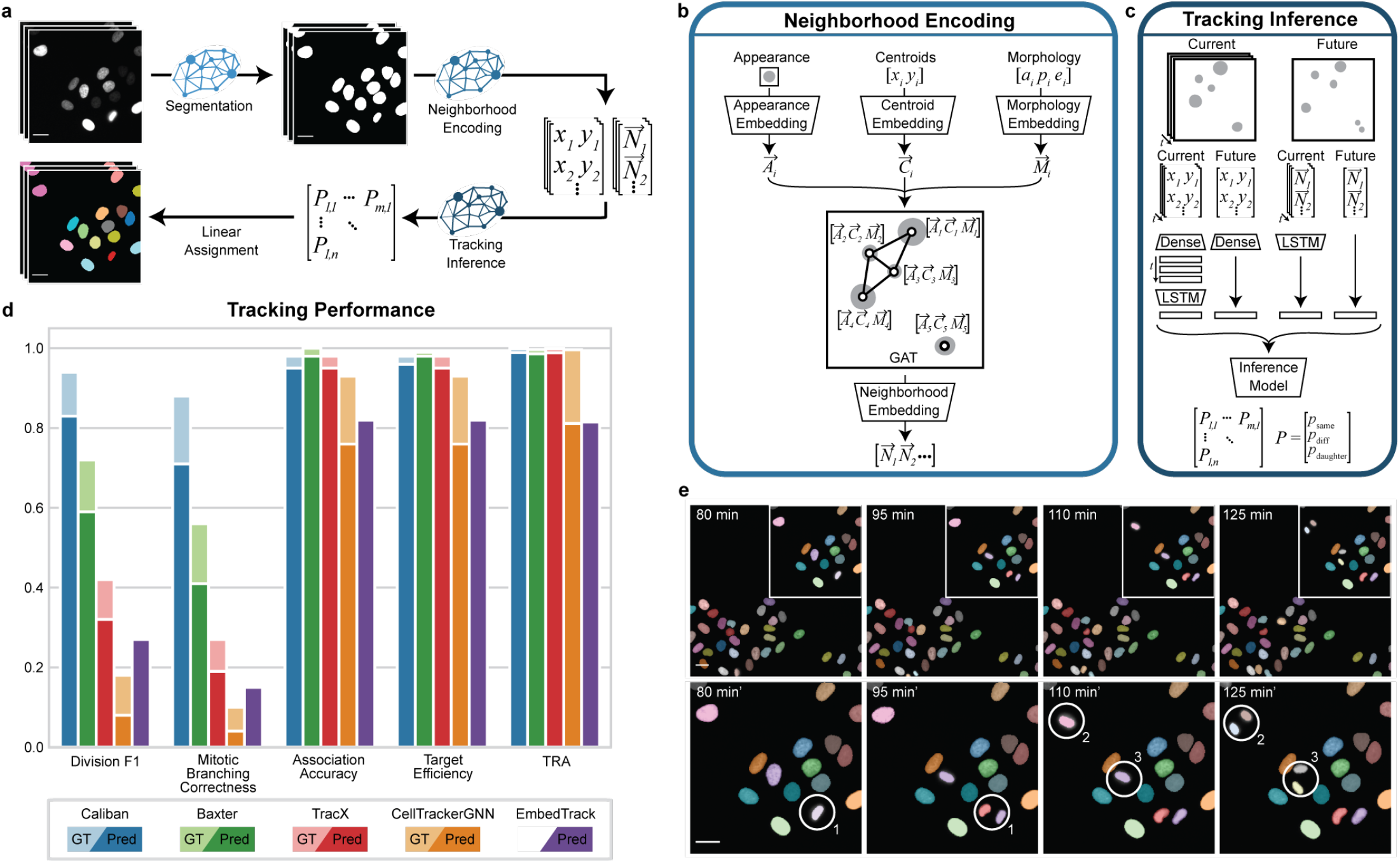
A deep learning approach to cell segmentation and tracking using Caliban. (a) Caliban takes a movie of fluorescently-labeled nuclei as input and then generates a nuclear segmentation mask for each frame. Features for each cell in a frame are extracted and passed through a neighborhood encoder model to generate a vector embedding for each cell. These embeddings and cell positions are passed into the tracking inference model, which predicts the probability that each pair of cells between frames is the same, is different, or has a parent–child relationship. These probabilities are used as weights for linear assignment to construct cell lineages on a frame-by-frame basis. (b) The neighborhood encoding model takes as input an image of each cell, its centroid position, and three metrics of morphology (area, perimeter, and eccentricity). A vector embedding of each input is used as node weights in a graph attention network^39,40^, where edges are assigned to cells within 64 pixels (41.6 *µ*m) of each other. The final neighborhood embedding for each cell captures the appearance of that cell and its spatial relationship with its neighbors in that frame. (c) The tracking inference model performs pairwise predictions on cells in frame *t*_*n*_ to cells in frame *t*_*n*+1_. The model is given neighborhood embeddings and centroid positions of cells in the previous seven frames [*t*_*n*−7_, *t*_*n*_] to compare with cells in frame *t*_*n*+1_. The temporal context of the previous seven frames is modeled using long short-term memory (LSTM) layers^41^. Ultimately, the model outputs a set of effective probabilities (*p*_same_, *p*_diff_, and *p*_parent-child_) for each pair of cells between frame *t*_*n*_ and frame *t*_*n*+1_. (d) The performance of Caliban and that of four other tracking methods were evaluated on the test split of DynamicNuclearNet. Tracking performance on ground-truth segmentations is excluded for EmbedTrack because it is an end-to-end method that generates segmentations as a part of tracking. TRA: tracking accuracy in the Cell Tracking Challenge. (e) A sample montage from DynamicNuclearNet with predictions from Caliban. Circles highlight the correct identification of three division events. (Scale bars = 26 *µ*m)

Separating our tracking model into two pieces facilitates rapid and scalable inference. During inference, the computationally expensive neighborhood encoder model can be run on all frames in parallel, leveraging GPU acceleration, followed by the lightweight tracking inference model, which is run on a frame-by-frame basis. The tracking inference model assigns lineages to cells by comparing the feature vectors of the last frame of existing lineages with the feature vectors of candidate cells in the current frame; model predictions are used with the Hungarian algorithm^44,45^ to complete the assignment. To accommodate the entry and exit of cells in the linear assignment framework, we create a “shadow object” for each cell in a frame, which allows assignments for the “birth” or “death” of cells^45^. (See Methods for details.)

We compared Caliban to four alternative methods that performed well in the Cell Tracking Challenge and could be run without manual parameter tuning — Baxter, CellTrackerGNN, EmbedTrack, and Trac^x^ — on a variety of metrics^33,46–48^. For each of these methods, we used pre-existing models or parameters that were trained or optimized on the Fluo-N2DL-HeLa dataset if available. For Caliban, we used a single version of the model that was trained on all five cell types represented in DynamicNuclearNet, including the two Fluo-N2DL-HeLa training movies. We tested each algorithm on ground truth and predicted segmentations. Predicted segmentations for each method were generated with that method’s segmentation model or Caliban’s segmentation model if the former was unavailable.

On measures of division performance evaluated on the DynamicNuclearNet testing split, Caliban outperformed all previously published methods. This performance boost is primarily attributable to Caliban’s cell-tracking capability rather than cell segmentation (Figure 2e), as the boost is present when tracking is performed on ground truth segmentations. On metrics focused on segmentation and linkages evaluated on the DynamicNuclearNet testing split, Caliban performed comparably to existing methods. For all metrics evaluated on the Cell Tracking Challenge Fluo-N2DL-HeLa test split, Caliban outperformed previously published methods (complete benchmarking results are shown in Supplementary Tables 2 and 3). We note that these benchmarks are unable to separate the relative contributions of training data size and model architecture to performance.

### Occident generates a catalog of phenotypes from segmented and tracked live-cell images

We applied the Caliban pipeline to segment and track individual cells from live-cell imaging of 1g4 anti-Ny-ESO-1 T cell receptor (TCR)-transgenic T cells (TCR T cells) co-cultured with cancer cells expressing NY-ESO-1 antigen (Figure 3a). Four replicates of three lines of TCR T cells, RASA2 KO, CUL5 KO, and SH KO T cells, at 1:1 ratio to A375 mKate^+^ tumor cells were collected. RASA2 KO and CUL5 KO T cells are known to demonstrate improved anti-cancer efficacy *in vitro* and are expected to show differences in behavior from the SH KO cells. Brightfield images, capturing T cells and cancer cells, and red fluorescent protein channel images, capturing cancer nuclei, were collected every four minutes over twenty-four hours. We applied Caliban segmentation with to identify putative T cells, cancer cells aggregates, and cancer nuclei, and then used Caliban’s tracking to link cell identifies across frames.

**Figure 3.**
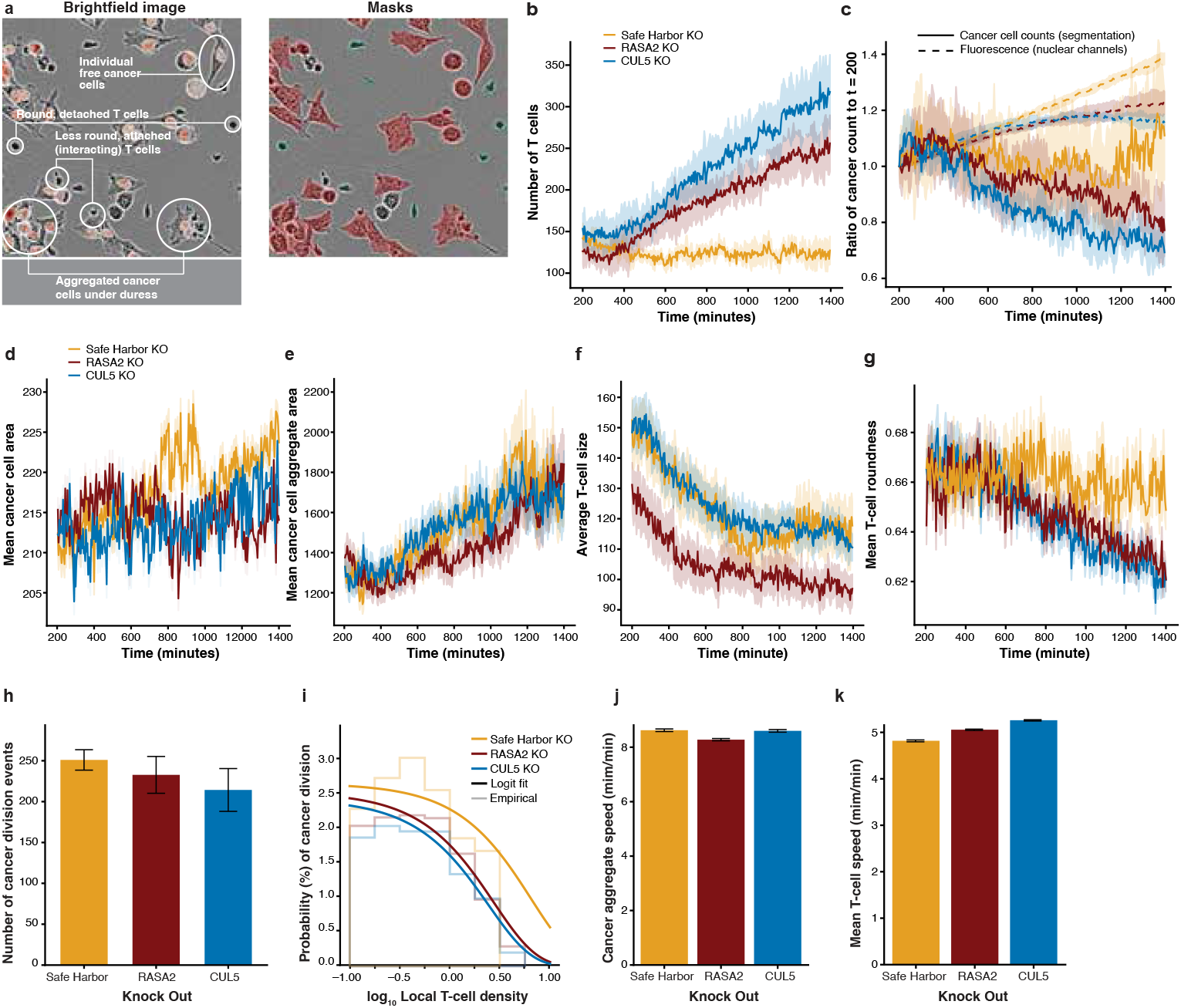
Extracting a catalog of cellular phenotypes with Occident. (a) Morphology is an indicator of cell state. T cells that are interacting with cancer cells change from their round shape to attack cancer cells while cancer cells aggregate together under distress. Segmentation masks are able to capture putative cells and cell agregates for both cell types. (b) Increased T cell expansion with beneficial knockouts. (c) Cancer cell population from segmentation over time reveals greater killing in wells with beneficial T cell perturbations. Fluorescence of the cancer nuclear marker is a lagging indicator of cancer cell population. (d) Cancer cells in RASA2 KO wells have lower average area, indicating greater stress from T cell attack. (e) SH KO and CUL5 KO T cells are smaller when attached to cancer cells, while RASA2 KO T cells increase in area when attached, likely due to greater activation. (f) The ratio of mean cancer cell clump area to mean individual cancer cell area increases over time as stressed cancer cells aggregate. (g) T cells with beneficial genetic knockouts become less round over time, indicating greater anti-cancer activity. (h) Cancer cell division events decreased in wells with T cells with beneficial genetic knockouts. (i) Increased local T cell density decreases the probability of cancer cell division and growth. T cells with beneficial genetic knockouts have a greater inhibitory effect on cancer cell division. (j) RASA2 KO T cells substantially decreased the average speed of cancer cells, suggesting greater anti-cancer effect during interactions. (k) T cells with beneficial genetic knockouts had higher average speed than control T cells, suggesting greater activity.

Segmentation allows us to interrogate the population of T cells and cancer cells. RASA2 or CUL5 knockouts in T cells are known to increase proliferation under suppressive conditions^7,13^. We observed that the number of CUL5 KO or RASA2 KO T cells in cancer co-culture wells increases over time, while the number of SH KO T cells remained approximately constant over time (Figure 3b). An exponential model of growth to quantify proliferation rates confirmed that CUL5 KO T cells (7.4 × 10^−4^ min^−1^, *p* ≤ 2.2 × 10^−16^) and RASA2 KO T cells (6.9 × 10^−4^ min^−1^, *p* ≤ 2.2 × 10^−16^) had substantially faster proliferation rates than SH KO T cells (−4.6 × 10^−5^ min^−1^).

RASA2 KO and CUL5 KO T cells are known to improve T cell’s *in vitro* cancer cell killing ability^7,13^. We observed a decrease in the number of cancer cells over time in those conditions relative to the SH KO condition (Figure 3c). The slope of a linear model of cancer cells over time revealed a substantial change in the rate of cancer killing activity for RASA2 KO T cells (− 2.8 ×10^−2^ min^−1^, *p* ≤ 2.2 × 10^−16^) and CUL5 KO T cells (−4.0×10^−2^ min^−1^, *p* ≤ 2.2 × 10^−16^) relative to SH KO T cells (−4.2 ×10^−3^ min^−1^). The number of cancer cells started to decrease before the fluorescent signal marking cancer cell nuclei levels off and decreases, suggesting that fluorescence of the nuclear reporter is a lagging signal of the cancer population (Figure 3c). The ordering of fluorescent intensity and number of cancer cells from segmentation was maintained; by both measures, cancer cells expanded the most when co-cultured with SH KO, followed by co-culture with RASA2 KO T cells. CUL5 KO best controlled the expansion of co-cultured cancer cells.

Occident characterized cell morphology across the genetic perturbations from cell segmentation. We prompted Segment Anything^49^ with nuclear masks to identify a subset of individual cancer cells composing the aggregates (Figure S5; see Methods for details). From the putative individual cancer cells identified, we observed that the mean area of these cells was consistent across time and the type of T cell co-cultured (Figure 3d). However, from the full cancer cells masks, we observed that aggregates increase in size over time (Figure 3e) across conditions. On average, we observed that aggregated cancer cells in co-culture with RASA2 KO T cells have lower mean area than with SH KO T cells (*p* ≤ 2.2 × 10^−16^), possibly indicating greater anti-cancer activity of this condition (Figure 3d). Interestingly, we observed that T cell size decreases over time across all three conditions (Figure 3). RASA2 KO T cells were on average smaller than CUL5 KO T cells (*p* ≤ 2.2 × 10^−16^) and SH KO T cells (*p* ≤ 2.2 × 10^−16^).

Roundness, the normalized ratio of area to perimeter that quantifies how similar a shape is to a perfect circle, is also a useful indicator of T cell. Free floating T cells demonstrate circular shapes (i.e., roundness close to 1), while attached T cells interacting with cancer cells are expected to change conformation, moving away from perfect roundness (Figure 3a). We observed that, as quantified by the slope of a linear regression fit, T cells with RASA2 KO (−1.4 × 10^−4^, *p* ≤ 2.2 × 10^−16^) and CUL5 KO (−1.8 × 10^−4^, *p* ≤ 2.2 × 10^−16^) become substantially less round over time compared to the SH KO control (−2.6 × 10^−5^ min^−1^; Figure 3g). Since decreased roundness is a hallmark of T cells interacting with cancer cells, the decrease in mean roundness over time suggests an increased number of T cells interacting with cancer cells in the genetic knockout conditions relative to control.

Occident takes advantage of tracking to reveal the temporal dynamics of cellular behavior. Cancer cell divisions are identified from cancer nuclei tracks. The difference in cancer divisions across perturbations was not statistically significant (Figure 3h). However, a logistic regression model to predict the probability of cancer division as a function of average local T count shows that an increase in the local T cell density is associated with decreased probability of a cancer division event as expected, due to the modified T cell’s anti-cancer cell function (Figure 3i). The decrease in log odds of a cancer cell division event per additional T cell is greater for the RASA2 KO T cells (− 0.37 events/cell^−1^, *p* ≤ 4 × 10^−3^) and CUL5 KO T cells (− 0.42 events/cell^−1^, *p* ≤ 7 × 10^−4^) relative to the SH KO T cells (− 0.16 events/cell^−1^; Figure 3i), again quantifying the stronger anti-cancer cell activity of these edited TCR T cells.

Occident also uses cell tracks to measure T cell and cancer cell speed, or the change in cell centroid position over time, to study cellular motility across genetic knockouts. We find that cancer cells co-cultured with RASA2 KO T cells had slower average speed (8.28 *µ*m/min; *p* ≤ 2.2 × 10^−16^) than cancer cells with SH KO T cells (8.62 *µ*m/min). Cancer cells in wells with CUL5 KO T cells (8.60 *µ*m/min; *p* ≤ 0.5) had similar speed to SH KO conditions (Figure 3j). The RASA2 KO (5.06 *µ*m/min, *p* ≤ 2.2 ×10^−16^) and CUL5 KO (5.26 *µ*m/min, *p* ≤ 2.2 × 10^−16^) T cells both had higher average speed than the SH KO T cells (4.82 *µ*m/min), suggesting greater T cell movement and activity (Figure 3k). Slower cancer cells suggest more debilitating effects from T cell attack from RASA2 KO cells, while faster T cells suggests improved hunting activity of the T cells with beneficial genetic perturbations^26^. Overall, RASA2 KO and CUL5 KO T cells had similar effects on cancer division, T cell velocity, and T cell roundness, but RASA2 KO T cells lead to greater changes in cancer cell size and speed. These differences in cell behavior phenotypes begin to paint the picture of the different mechanism-of-actions between genetic perturbations that would be missed without dynamic imaging quantification and behavioral phenotyping.

### Occident characterizes cellular changes after T cell-cancer cell interaction

Occident characterizes the cellular behavior phenotypes during T cell-cancer cell interactions to understand cellular behavior, revealing changes across the three conditions. An *interaction* between a T cell and a cancer cell is defined as at least two consecutive frames with overlapping masks between the two cells (Figure 4a). We observed that wells with RASA2 KO T cells (*p* ≤ 2.2 ×10^−16^, Wilcoxon signed-rank test) and CUL5 KO T cells (*p* ≤ .001) show more interactions over time than wells with SH KO T cells (Figure 4b).

**Figure 4.**
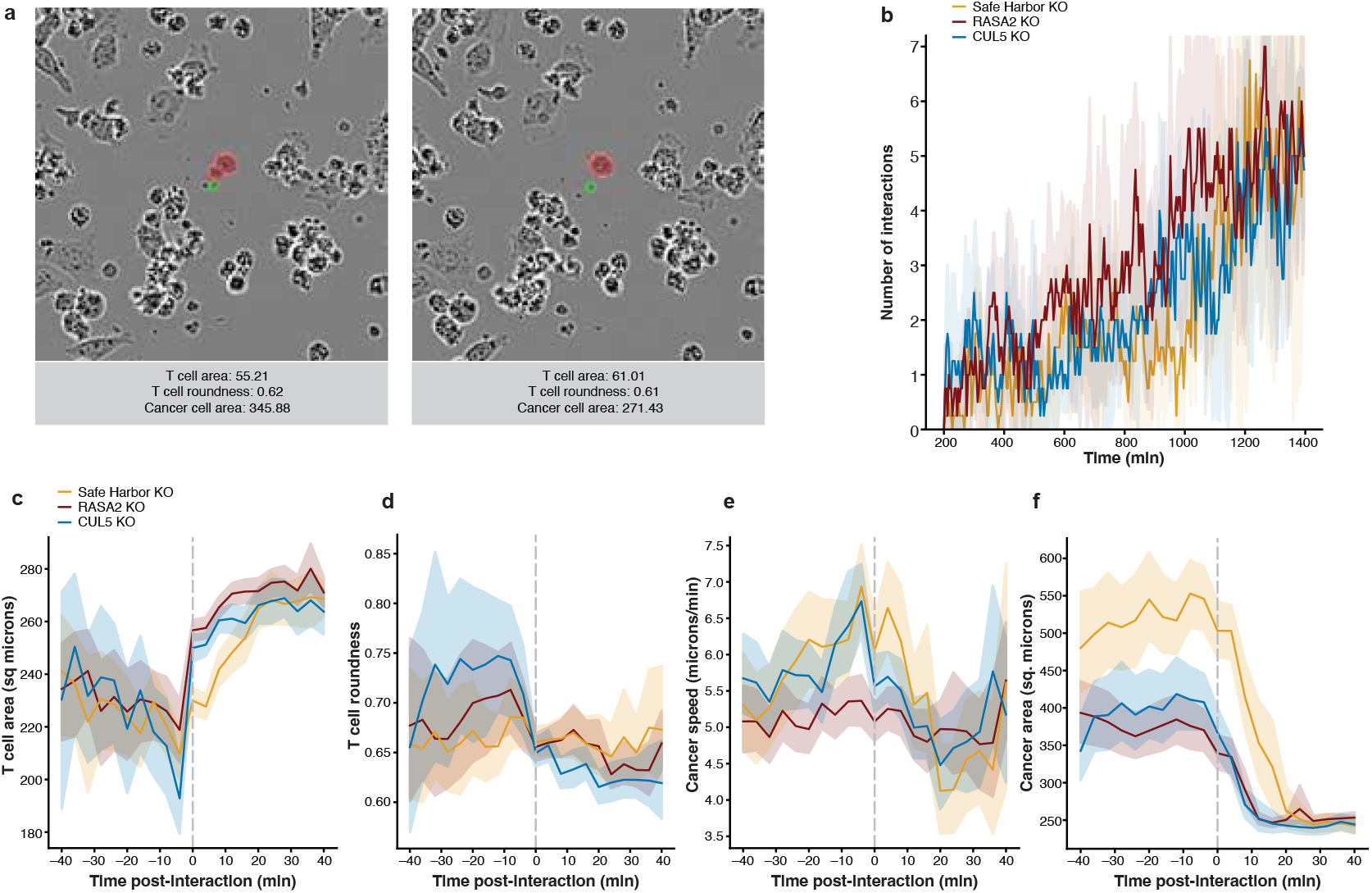
Occident characterizes the behavior of T cells and cancer cell aggregates during cell-cell interactions. (a) An example interaction between a SH KO T cell and a cancer cell, and the same cells 4 minutes later. (b) Wells with RASA2 KO T cells have a higher average number of cancer cell – T cell interactions across time than CUL5 KO and SH KO wells. (c) Average T cell size increased after interaction across conditions. (d), T cell roundness decreased as they interact with cancer cells. (e) Cancer aggregate speed decreased during interactions with T cells. (k) Cancer cells decreased in area after interaction with T cells.

Following the phenotypes of interacting cells reveals complex dynamics of T cell–cancer cell interactions. T cells have been observed to increase in size after activation from interacting with an antigen, such as a cancer cell^50,51^. We indeed found that T cell area increased after interaction with cancer cells across all three conditions (Figure 4c). For example, SH KO T cells increased from 225.8 *µ*m^2^ average area for the forty minutes pre interaction with cancer cells to 241.7 *µ*m^2^(*p* ≤ 2.2 × 10^−16^) eight minutes post interaction and 268.6 *µ*m^2^(*p* ≤ 2.2 × 10^−16^) forty minutes post interaction. The increase in average area of RASA2 KO and CUL5 KO T cells was similar. T cells become less round after interaction with cancer cells (Figure 4d); CUL5 KO T cells show a marked drop from an average roundness of 0.65 for the forty minutes pre interaction to 0.61 (*p* ≤ 2.2 × 10^−16^) eight minutes post interaction, increasing slightly to 0.62 (*p* ≤ 2.2 × 10^−16^) forty minutes post interaction. Taken together, these results allow us to quantify and optimize T cells for activation after interacting with antigens.

Interactions with T cells also affect cancer aggregates. Cancer cell speed decreases post interaction in all three conditions (Figure 4e). When co-cultured with CUL5 KO T cells, for example, cancer cells slow from 5.88 *µ*m/min (*p* ≤ 2.2 × 10^−16^) on average for forty minutes pre interaction to 5.51 *µ*m/min (*p* ≤ 2.2 × 10^−16^) eight minutes post interaction, dipping all the way down to 4.47 *µ*m/min (*p* ≤ 2.2 ×10^−16^) twenty minutes post interaction, then rebounding to 5.18 *µ*m/min (*p* ≤ 2.2 × 10^−16^) forty minutes post interaction. Restricting analysis to masks with putatively single-cell level areas, we also observe a decrease in area post-interaction across all three conditions (Figure 4f). All three conditions reduce the mean cancer area to about 250 sq. *µ*m. Cancer cells in RASA2 KO and CUL5 KO wells were already smaller on average to start with, likely from previous interactions with T cells. We visually observe an increase in T cell area, a decrease in T cell roundness, and a decrease in cancer cell area four minutes post interaction (Figure 4b).

### Spatiotemporal modeling reveals changes in T cell inter-cellular signaling and proliferation under genetic perturbation

Anti-cancer activity of T cells results from group behavior. To quantify multi-cellular spatiotemporal dynamics, we estimated the *Markov transition* matrix of local cell density states. Briefly, we split each video spatially into non-overlapping windows or bins, and we define each bin’s state at each time point as the number of T cells and number of cancer cells within the bin. The Markov transition matrix is estimated by observing how often a bin in state *τ* T cells, 𝒷 cancer cells at time *t* moves to state (*τ*^′^, 𝒷 ^′^) at time *t* + 1 (Figure 5a).

**Figure 5.**
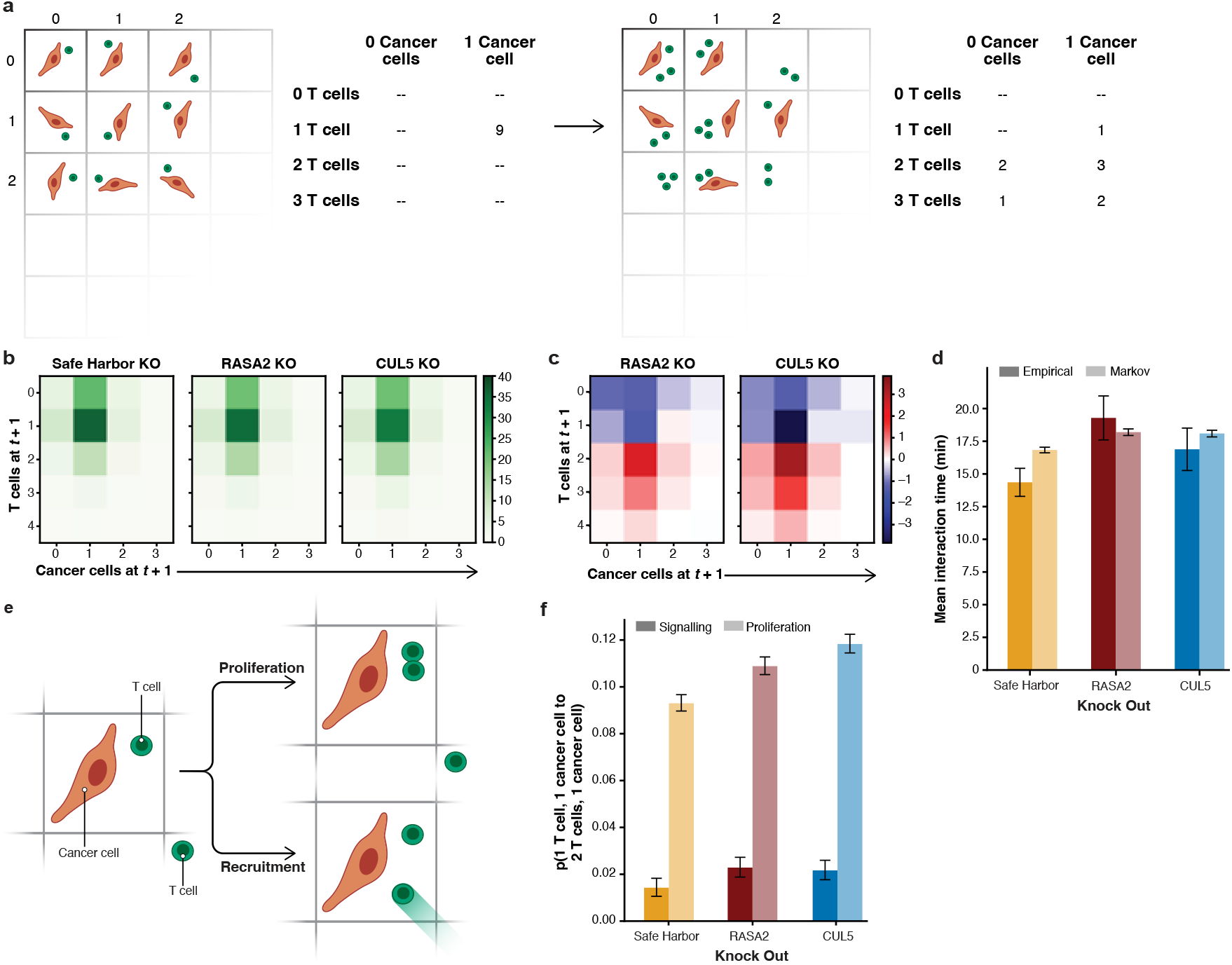
An interpretable spatiotemporal model of cell organization characterizes interaction with cancer cell duration, recruitment, and proliferation in T cells. **a**, Schematic of Markov transition model and identification of proliferation and recruitment events. **b**, Estimated Markov transition probabilities of states at *t* + 1 from 1 T cell, 1 cancer cell at time *t* by genetic perturbation **c**, difference in Markov transition probabilities at *t* + 1 from 1 T cell, 1 cancer cell at time *t* from SH KO control. **d** Mean time of interaction estimated from tracks and Markov transition probabilities, controlling for two frame (eight minute) minimum cutoff. **e**, Probability of transition from 1 T cell, 1 cancer cell at time *t* to 2 T cells, 1 cancer cell at *t* + 1 from recruitment and from proliferation, by genetic perturbation.

The Markov transition matrix captures the temporal dynamics of cell densities between frames. To understand how individual T cell–cancer cell interactions lead to multi-cellular behavior, we focus on the initial state of *τ* = 1 T cell and 𝒷 = 1 cancer cell. We find that the *1 T cell, 1 cancer cell* state is most likely to remain the same in the next observation (Figure 5b) in all three conditions. We use the probability of detachment, defined as the likelihood of transition from *1 T cell, 1 cancer cell* to *zero T cells and any number of cancer cells*, to estimate the duration of interaction (or *dwell time*) with a negative binomial model (see Methods for details). Control SH KO T cells had a shorter dwell time (16.83 ± 0.21 min) compared to RASA2 KO T cells (18.20 ± 0.25 min, *p* ≤ 5.1 ×10^−16^) and CUL5 KO T cells (18.09 ± 0.25 min, *p* ≤ 1.2 × 10^−13^). This matches the results of the empirical mean interaction times; compared to SH KO (14.35 ± 1.08 min), RASA2 KO (19.28 ± 1.68 min, *P* ≤ 1.9 × 10^−6^) and CUL5 KO (16.88 ± 1.62 min, *p* ≤ 9.7 ×10^−3^) have empirically longer interaction times with cancer cells (Figure 5d). These results present another indicator of the beneficial anti-tumor effects of the genetic perturbations. The average interaction times around 20 minutes are also visible in the cancer phenotypes post interaction (Figure 4k,l), where cancer cell properties start to return to the pre-interaction baseline 20 minutes post-interaction. These consistent results validate the Markov model as an effective estimator of cell dynamics.

Beyond the individual cell dynamics, T cells activated by interacting with antigens such as cancer cells may use molecular signals to *recruit* more T cells and *proliferate*^52^. RASA2 KO and CUL5 KO T cells have previously been observed to proliferate faster than SH KO T cells^7,13^ and increase the production of cytokines such as IL-2, IFN*γ*, and TNF^7,13^, a molecule involved in T cell recruitment. Comparing the transition probabilities of edited T cells to the SH KO T cells revealed that the RASA2 KO (23.63%, *p* ≤ 2.2 × 10^−16^) and CUL5 KO (26.42%, *p* ≤ 2.2 × 10^−16^) T cells were more likely to increase in number at the next time point from this state than SH KO T cells (18.64%).

To deconvolve the contributions of T cell recruitment versus proliferation to this transition, we quantify the transitions from *1 T cell, 1 cancer cell* to *2 T cells, 1 cancer cell* from tracked cells, indicating a recruitment event, versus untracked cells, indicating a T cell proliferation event (Figure 5e). We observed that RASA2 KO (2.30%; *p* ≤ 2.2 × 10^−16^) and CUL5 KO (2.18%; *p* ≤ 2.2 ×10^−16^) T cells have greater recruitment probability than SH KO T cells (1.44%), and RASA2 KO and CUL5 KO are similar in their recruitment probability (*p* ≤ 0.31; Figure 5f). CUL5 KO (11.85%, *p* ≤ 2.2 × 10^−16^) and RASA2 KO (10.90%, *p* ≤ 7.8 ×10^−11^) T cells have a higher probability of proliferation upon interaction than SH KO (9.32%), with CUL5 KO T cells more proliferative than RASA2 KO T cells (*p* ≤ 2 × 10^−4^). These results suggest that the CUL5 KO enhanced T cell proliferation, whereas RASA2 KO more notably increased T cell interaction time with cancer cells.

## Discussion

Live-cell imaging captures rich data on T cell and cancer cell morphology, motility, and behavioral dynamics. The combination of modern computational techniques and capacity with increased imaging resolution and throughput allows for cellular behavior analysis to push our understanding of complex multi-cellular behaviors forward. Improved quantification of population behavioral dynamics of cells will lead to better understanding of basic biology and direct the engineering and optimization of new precision treatments. In this paper, we first developed a state-of-the-art cell segmentation and tracking platform, Caliban, based on human-in-the-loop deep learning, to serve as the basis of cellular behavior analysis. We then developed the Occident pipeline to quantify morphological and behavioral phenotypes in co-cultures of TCR T cells and cancer cells. Caliban’s improved tracking combined with Occident’s interpretable modeling revealed biologically-important differences between the beneficial *RASA2* and *CUL5* knockouts. In particular, RASA2 KO T cells spent more time attached to cancer cells, decreasing cancer speed and size, while CUL5 KO T cells proliferated more upon activation by cancer cell interactions, creating more T cells to fight tumor cells.

This cellular behavioral analysis of RASA2 KO and CUL5 KO T cells is a first step to demonstrate the utility of these techniques. Challenges remain in terms of segmentation and tracking quality. Many cells remain unidentified or are characterized by imprecise masks. Many of the segmented T cells that do not move may be larger detritus in the well rather than a T cell. The mismatch between cancer cell count and cancer cell fluorescence indicates there may be interesting behavior inside the cancer cell aggregates that segmentation struggles to capture well. Tracking is limited in its ability to detect fast moving cells, challenging our ability to quantify rapidly dynamic behaviors. One-step Markov transitions are a simple class of models that may miss longer interval behavior or more localized spatial behaviors. Wells are imaged from above in 2D, masking z-axis behaviors, while cancer cells regularly adhere to the well walls, elongating their shape; these experimental conditions and their effects on the data must be explored. Better image processing will enable richer models that can more clearly identify detachment, proliferation, and recruitment, and extend to phenotypes such as T cell swarming and tracking.

Despite challenges, Caliban and Occident quantify complex and therapeutically-relevant cellular morphology and behavior phenotypes with currently available common laboratory research hardware and software, capturing important signals from these ubiquitous live-cell imaging data. Further improvements in imaging techniques, e.g., capturing additional reporters, three dimensions, or greater temporal and spatial resolution, will further enable a deeper investigation of spatial behavior. As more data are collected from ever-improving screening techniques, consisting of more diverse cell types, donors, and perturbations, among other variables, more impactful insights in the changes to morphology, proliferation, signaling, and activation behaviors induced by the cell types, perturbations, and interactions will be uncovered. Looking forward, we expect these insights can be combined with emerging genomics foundation models to predict genetic modifications that program desired cellular behaviors. Our work opens the door to directly quantifying and analyzing dynamic multicellular behavior phenotypes captured with live-cell imaging to predict, design, and optimize cell therapies.

## Methods

### Caliban data collection

#### Cell culture

We used five mammalian cell lines (NIH-3T3, HeLa-S3, HEK293, RAW 264.7, and PC-3) to collect training data. All lines were acquired from the American Type Culture Collection. We cultured the cells in Dulbecco’s modified Eagle’s medium (DMEM; Invitrogen; RAW 264.7, HEK293, and NIH-3T3) or F-12K medium (Caisson; Hela-S3 and PC-3) supplemented with 2 mM L-glutamine (Gibco), 100 U/mL penicillin, 100 *µ*g/ml streptomycin (Gibco), and either 10% calf serum (Colorado Serum Company) for NIH-3T3 cells or 10% fetal bovine serum (FBS; Gibco) for all other cells.

#### Live-cell imaging

Before imaging, cells were seeded in fibronectin-coated (10 *µ*g/mL; Gibco) glass 96-well plates (Nunc or Cellvis) and allowed to attach overnight. We performed nuclear labeling via prior transduction with H2B-iRFP670 (Hela, RAW 264.7), H2B-mClover (HEK293, NIH/3T3), and H2B-mCherry (PC-3). The media was removed and replaced with imaging media (FluoroBrite DMEM (Invitrogen) supplemented with 10 mM HEPES (Sigma-Aldrich), 10% FBS (Gibco), 2mM L-glutamine (Gibco)) at least 1 h before imaging. We imaged cells with a Nikon Ti-E or Nikon Ti2 fluorescence microscope with environmental control (37^*°*^C, 5% CO_2_) and controlled by Micro-Manager or Nikon Elements. We acquired images at 5-to 6-min intervals with a 20x objective (40x for RAW 264.7 cells) and either an Andor Neo 5.5 CMOS camera with 3 ×3 binning or a Photometrics Prime 95B CMOS camera with 2 ×2 binning. All data were scaled so that pixels had the same physical dimensions (0.65 *µ*m per pixel) before training.

#### Caliban dataset development: DeepCell Label

We previously described DeepCell Label^38^, our browser-based software for data annotation. We extended DeepCell Label to support labeling cell lineages and divisions in dynamic datasets. Additionally, we implemented a state machine that allows annotators to apply undo/redo functions during their work. These new features are described below.

DeepCell Label manages the state of its labeled data with a Python-based backend and a React-based frontend. The backend serves and submits project data and provides image processing for editing label arrays, while the frontend controls the user interface and edits non-image-based labels. The front end retrieves images and labels for the project from the backend and loads the data into its state management.

We use the Javascript library XState to manage the state on the front end. Through XState, we define actors that manage the state of both user interface (UI) elements and data labels. An actor consists of a context, containing arbitrary data, along with a set of states and transitions between states. Actors receive events from other actors or the user, and each state defines how to transition between states and update its context upon receiving events. Some actors maintain UI state, while others define and control the operations that can edit labeled data. The application creates a root actor that instantiates a tree of child actors corresponding to each UI element or type of labeled data. The root actor sets up communication between child actors, enabling features like undo/redo that must orchestrate state across the application.

Actors expose their state to user interface elements by React’s hooks, allowing multiple components to access and update the same shared state. For instance, to adjust the contrast of an image, a Slider can expose the contrast settings for a user to update, while a Canvas component can access the updated contrast settings to render the image. For example, we define an actor for the image canvas that handles mouse movement events with context-dependent behavior. When panning around the canvas, a mouse-move event changes the position of the canvas, while when not panning, we update the coordinates for the cursor. Clicking and releasing the mouse sends mouse-up and mouse-down events, transitioning to and from the panning state.

We implement our undo feature via an undo actor that records and restores the states of both UI and label actors over time. The undo actor maintains two stacks of project states: an undo stack with past snapshots to restore and a redo stack with undone snapshots. Editing the labels after undoing clears the redo stack, so only one branch of the project state is maintained. To integrate with the undo feature, actors register themselves with the undo actor and agree to submit snapshots of their state that can be restored upon undoing or redoing an action. Two types of actors can register with the undo actor: UI actors that manage the state of UI elements, and label actors that manage and edit labels. When a label actor edits its labels, it submits a SNAPSHOT event to the undo actor containing a copy of the labels before and after editing. The undo machine then broadcasts a SAVE event to all UI actors, which respond with a RESTORE event containing their current state. When the user undoes an action, the SNAPSHOT and RESTORE events are resent to the actors that originally submitted the events. With this approach, each UI and label actor is responsible for defining how its state should be recorded and restored with the events it submits, and the undo actor is responsible only for orchestrating and broadcasting the events to all registered actors. As new types of labeled data and UI elements are developed, the actors that drive new features can flexibly integrate with the undo infrastructure by defining these events for themselves and registering with the undo actor.

#### Data annotation

In this study, we used DeepCell Label in two stages to generate a nuclear tracking dataset. First, annotators were asked to correct nuclear segmentation labels for all frames in the dataset. Movies were broken into five frame sets for segmentation which allows annotators to leverage the temporal context present in the movie to improve the annotation of dividing cells. Second, after segmentation annotations were complete, annotators were asked to label the nuclear segmentation masks such that a single cell maintains the same label across frames. Additionally, all division events were annotated with the connection of the parent cell to each daughter cell. An expert annotator reviewed all annotated movies before incorporation into the training dataset. The supplementary information provides a user manual for DeepCell Label (Supplementary File 1), along with sample instructions for segmentation (Supplementary File 2) and tracking (Supplementary File 3) corrections. Annotations were conducted by a team of four annotators and two expert reviewers. Each movie was annotated by a single annotator and approved by a single expert eliminating the need for resolving differences between two independent annotators.

#### Data versioning with DVC

Each labeled movie was versioned and tracked with DVC^53^. We recorded additional metadata in each `.dvc` file, including the data dimensions, annotation progress, and data source. These metadata enabled automatic data processing for generating segmentation and tracking predictions as well as launching annotation tasks.

#### Dataset sources

DynamicNuclearNet contains data from six sources. Five datasets were collected internally as described in sections Cell Culture and Live Imaging. Additionally, the CTC dataset Fluo-N2DL-HeLa was incorporated after generating complete segmentation masks for all frames using the protocol described above.

### Caliban Nuclear segmentation

#### Deep learning model architecture

The deep learning model for nuclear segmentation was based on the design of feature pyramid networks^54,55^. The network was constructed from an EfficientNetV2L backbone^56^ connected to a feature pyramid. Input images were concatenated with a coordinate map before entering the backbone. We used backbone layers C1–C5 and pyramid layers P1–P7. The final pyramid layers were connected to three semantic segmentation heads that predict transforms of the labeled image.

#### Label image transforms

For each image, we used a deep learning model to predict three different transforms, as inspired by previous work^38,57,58^. The first transform predicted whether a pixel belongs to the foreground or background, known as the “foreground–background transform.” The second transform predicted the distance of each pixel in a cell to the center of the cell and is called the “inner distance.” If the distance between a pixel and the center of the cell is *r*, then we compute the transform as 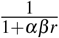, where 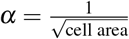 and *β* is a hyperparameter set to 1_38_. The final transform was the “outer distance,” which is the Euclidean distance transform of the labeled image. The loss function was computed as the sum of the mean squared error on the inner and outer distance transforms and the weighted categorical cross-entropy^59^ on the foreground–background transform. The cross-entropy term was scaled by 0.01 before the sum.

#### Preprocessing

Each image was required to have a minimum of one labeled object. Additionally, each image was normalized using contrast-limited adaptive histogram equalization with a kernel size equal to 1/8 of the image size to ensure that all images have the same dynamic range^60^.

#### Postprocessing

We fed two of the three model outputs, the inner and outer distance, into a marker-based watershed method^61^ to convert the continuous model outputs into a discrete labeled image in which each cell is assigned a unique integer. We applied a peak-finding algorithm^62^ with a radius of 10 pixels and a threshold of 0.1 to the inner distance prediction to determine the centroid location of each cell. Next, we generated the cell mask image by applying the watershed algorithm to the inverse outer distance prediction with the centroids as markers and a threshold of 0.01.

#### Model training and optimization

Training data were augmented with random rotations, crops, flips, and scaling to improve the diversity of the data. We used 70% of the data for training, 20% for validation, and 10% for testing. The model was trained using the Adam optimizer^63^ with a learning rate of 10^−4^, a clipnorm of 10^−3^, and a batch size of sixteen images; training was performed for sixteen epochs. After each epoch, the learning rate was adjusted using the function lr = lr ×0.99^epoch^. Additionally, if the loss of the validation data did not improve by more than 10^−3^ after five epochs, the learning rate was reduced by a factor of 0.01.

To optimize the model’s performance on nuclear segmentation, we tested ten backbones: ResNet50^64^, ResNet101^64^, EfficientNetB2^65^, EfficientNetB3^65^, EfficientNetB4^65^, EfficientNetV2M^56^, EfficientNetV2L^56^, EfficientNetV2B1^56^, Efficient-NetV2B2^56^, and EfficientNetV2B3^56^. Additionally, we explored the optimal set of pyramid layers: P1–P7 and P2–P7.

#### Evaluation

To fully evaluate the performance of our segmentation model, we developed a set of object-based error classes that assess the model on a per-object basis as opposed to a per-pixel basis. This framework provided a perspective on model performance that reflects downstream applications. First, we built a cost matrix between cells in the ground truth and cells in the prediction, where the cost is one minus the intersection over union (IoU) for each pair of cells. We performed a linear sum assignment on this cost matrix, with a cost of 0.4 for unassigned cells, to determine which cells were correctly matched between the ground truth and prediction. For all remaining cells, we constructed a graph in which an edge was established between a ground truth and a predicted cell if the IoU was greater than zero. For each subgraph, we classified the error type based on the connectivity of the graph. Nodes without edges corresponded to a false positive or negative if the graph contained only a predicted or ground truth cell, respectively (Supplementary Fig. 2a–c). A single predicted node connected to multiple ground truth nodes indicated a merge error (Supplementary Fig. 2d). Conversely, a single ground truth node connected to multiple predicted nodes was a split error (Supplementary Fig. 2e). Finally, any subgraphs that contain multiple ground truth and predicted nodes were categorized as “catastrophe” (Supplementary Fig. 2f). The resulting error classes can be used to calculate a set of summary statistics, including recall, precision, and F1 score by using the true positive, false positive, and false negative classes. The remaining error classes can be used to calculate (1) the number of missed detections resulting from a merge, (2) the number of gained detections resulting from a split, (3) the number of true detections involved in a catastrophe, and (4) the number of predicted detections involved in a catastrophe.

### Caliban cell tracking

#### Linear assignment for tracking

Tracking was treated as a linear assignment problem^45^. To solve the tracking problem, we first constructed a cost function for possible pairings across frames. The tracking problem was then reduced to the selection of one assignment out of the set of all possible assignments that minimized the cost function. This task can be accomplished with the Hungarian algorithm^66^. One complicating factor of biological object tracking is that objects can appear and disappear, which leads to an unbalanced assignment problem. Cells can disappear by either moving out of the FOV or dying. Similarly, cells can appear by moving into the FOV or dividing into two daughter cells from one parent cell. In the context of the linear assignment problem, one can preserve the runtime and performance by introducing a “shadow object” for each object in the two frames that represents an opportunity for objects to “disappear” (if an object in frame *t*_*n*_ is matched with its shadow object in frame *t*_*n*+1_) or “appear” (if an object in frame *t*_*n*+1_ is matched with its shadow object in frame *t*_*n*_)^45^. Assuming that mitotic events can be accommodated by a “shadow object” as well, division detection and assignment fit neatly into this framework. This framework can also accommodate cells that disappear from the field of view and reappear, by allowing unmatched cells from prior frames that were not assigned to cell division events to participate in the assignment. With the annotated trajectories and divisions from our dataset, it then becomes a matter of developing a deep learning architecture to extract an object’s features and learn an optimal cost function.

To construct our learned cost function, we cast it as a classification task. Let us suppose that we have two cells: our target cell *i* in frame *t*_*n*_ and cell *j* in frame *t*_*n*+1_. Our goal was to train a classifier that takes in information about each cell and produces an effective probability indicating whether these two instances are the same, are different or have a parent–child relationship. If we have already tracked several frames, we incorporate temporal information by using multiple frames of information for cell *i* as an input to the classifier. This approach allowed us access to temporal information beyond just the two frames we are comparing. Our classifier was a hybrid deep learning model that blends recurrent, convolutional, and graph submodels; its architecture is summarized in Fig. 2b,c. The three scores that the model outputs, (*p*_same_, *p*_diff_, and *p*_parent-child_), which are all positive and sum to unity, can be thought of as probabilities. These scores were used to construct the cost matrix. If a cell in frame *t*_*n*+1_ is assigned to a shadow cell, i.e., if it “appears,” then we assessed whether there is a parent–child relationship. This was done by finding the highest *p*_parent-child_ among all eligible cells (i.e., the cells in frame *t*_*n*_ that were assigned to “disappear”)—if this probability was above a threshold, then we made the lineage assignment.

#### Neighborhood encoder architecture

To capture the contextual information of each cell and its neighbors, we constructed a graph attention network^39,40^. There were three input heads to the model. The first head received images of each cell and converted these images to a vector embedding with a convolutional neural network. Each image consisted of a 16 ×16 crop of the raw data centered on the centroid position of the cell. Additionally, the pixels within the nuclear segmentation mask were normalized by subtracting the mean value and dividing by the standard deviation. The second head received the centroid location of each cell. The third head received three morphology metrics for each cell: area, perimeter, and eccentricity. The latter two heads made use of fully connected neural networks to convert the inputs into vectors. We built an adjacency matrix for the graph attention network based on the Euclidean distance between pairs of cells; cells were linked if they were closer than 64 pixels (41.6 *µ*m). The normalized adjacency matrix and concatenated embeddings were fed into a graph attention layer^39^ to update the embeddings for each cell. The appearance and morphology embeddings were concatenated to the output of the graph attention layer to generate the final neighborhood embedding.

#### Tracking model architecture

Given cell 1 in frame *t*_*n*_ and cell 2 in frame *t*_*n*+1_, the neighborhood encoder was used to generate embeddings for cell 1 in frame *t*_*n*_ and the previous seven frames [*t*_*n−*7_, *t*_*n*_]. Long short-term memory^41^ layers were applied to the resulting embedding for cell 1 to merge the temporal information and to create a final summary vector for cell 1. The neighborhood encoder then generated an embedding for cell 2. Next, the vectors for cell 1 and cell 2 were concatenated and fed into fully connected layers. The final layer applied the softmax transform to produce the final classification scores: *p*_same_, *p*_diff_, and *p*_parent-child_.

#### Training and optimization

Both the neighborhood encoder and the inference model were jointly trained end-to-end such that the neighborhood embedding was tuned for the inference task. The model was trained on data that compare a set of frames [*t*_*n−*7_, *t*_*n*_] with frame *t*_*n*+1_. Each comparison of *t*_*n*_ with *t*_*n*+1_ contributed to the loss. For inference, the model was given single pairs of frames, e.g., *t*_*n*_ vs. *t*_*n*+1_. Training data were augmented with random rotations and translations. We used 70% of the data for training, 20% for validation, and 10% for testing. Data splitting was performed with regard to the cell type such that each cell type is equally represented across the three splits. The model was trained using the rectified Adam optimizer^67^ with a learning rate of 10^−3^, a clipnorm of 10^−3^, and a batch size of eight images. After each epoch, the learning rate was adjusted using the function lr = lr ×0.99^epoch^. Additionally, if the loss of the validation data did not improve by more than 10^−4^ after five epochs, the learning rate was reduced by a factor of 0.1. The model was trained over 50 epochs.

To optimize the performance of the tracking model, we tested the following parameters: graph layers (graph convolution layer, graph convolution layer with trainable skip connections, and graph attention convolution layer), distance threshold (64, 128, 256 pixels; 41.6, 83.2, 166.4 *µ*m), crop mode (fixed and resized), birth probability, division probability, and death probability.

#### Evaluating tracking performance

To evaluate the tracking performance, we used two sets of metrics. The first set assessed the linkages between cells, whereas the second set focused on the linkages of dividing cells. For the first set of metrics, we calculated the target efficiency (TE) and association accuracy (AA)^47,48^. Briefly, TE assesses the fraction of cells assigned to the correct lineage, and AA measures the number of correct linkages generated between cells.

Traditional metrics for evaluating tracking, including TE and AA, do not accurately reflect the ability of the method to identify divisions because divisions are relatively rare events. To overcome this weakness, we developed an evaluation pipeline that classifies each division event as a correct, missed, or incorrect division. Our pipeline can handle tracking assignments on ground truth and predicted segmentations. First, we calculated the IoU between cells in the ground truth and the predictions to establish a mapping that can be used to compare tracking predictions. On predicted segmentations, an IoU threshold of 0.6 was used as a threshold for overlap. For each division in the ground truth, we checked the corresponding node in the prediction to determine whether it was labeled as a division. If the daughter nodes in the prediction match those in the ground truth, the division was counted as a correct division (Supplementary Fig. 3a). We have found that, depending on the predicted segmentations, a division can sometimes be assigned to the frame before or after the frame that is annotated as a division in the ground truth data. We treated these shifted divisions as correct or a true positive. If the predicted node was not labeled as a division, it was considered as a missed division or false negative (Supplementary Fig. 3b). Finally, if a predicted parent node was identified as a division, but the daughters did not match the ground truth daughters, the division was counted as incorrect and included as a false negative division (Supplementary Fig. 3c). Finally, any remaining predicted divisions that cannot be matched to a ground truth division are counted as false positives.

We used the classified divisions to calculate a set of summary statistics, including recall, precision, and F1 score. Additionally, we used the mitotic branching correctness (MBC) metric^33^, calculated as follows:

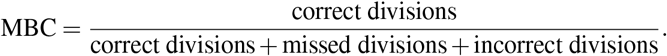

### Deployment

We previously described the DeepCell Kiosk^68^, our scalable cloud-based deployment for deep learning models. The Kiosk provides a drag-and-drop interface for model predictions currently deployed at www.deepcell.org/predict. To provide a seamless pipeline for nuclear segmentation and tracking, we deployed a new consumer for tracking jobs. First, each movie is split into single frames, which are distributed for nuclear segmentation. This step takes advantage of the Kiosk’s ability to parallelize and scale resources to match demand. Once nuclear segmentation is complete on all frames, the masks are concatenated, and tracking is performed. The user receives a final output that contains the raw data, labeled masks, and lineages.

### Benchmarking

We compared the performance of our model against four other algorithms: Baxter^69^, CellTrackerGNN^70^, EmbedTrack^71^, and Trac^x72^. Using the test split of our dataset, we evaluated the tracking performance of each algorithm on ground truth segmentation and predicted segmentations generated by either the algorithm or Caliban. We evaluated the resulting tracking predictions using our division evaluation pipeline and evaluation software from the Cell Tracking Challenge^46^. To evaluate the performance of Caliban on CTC movies, we manually annotated the two test movies from the Fluo-N2DL-HeLa dataset. To preserve the integrity of the challenge, we have not released our annotations but used them for evaluation in this paper. The notebooks used to generate benchmarks are available at https://github.com/vanvalenlab/Caliban-2024_Schwartz_et_al.

We evaluated Caliban’s inference speed using a single GPU (NVIDIA RTX A6000) and eight CPUs (AMD EPYC 7763 64-Core Processor). The inference time was split into four sections: segmentation inference, neighborhood encoder inference, tracking inference, and linear assignment. Inference was repeated three times for each movie in the test data split.

### Occident data collection

#### Collection and culture of primary T cells

Leukopaks from anonymized donors were acquired from StemCell Technologies with approved IRBs. Human primary T cells were then isolated with the EasySep Human T Cell Isolation Kit (StemCell Technologies). The T cells were cultured at a density of 1 million cells per ml maintained in X-Vivo-15 medium supplemented with 5% fetal bovine serum, 50 *µ*M beta-mercaptoethanol, and 10 mM N-acetyl-L-cysteine plus 100 IU/mL of IL-2. They were then activated with Dynabeads Human T-Activator CD3/CD28 (Gibco) at a 1:1 bead-to-cell ratio.

#### Lentiviral generation followed by T cell transduction for TCR expression

Lenti-X 293T cells (Takara Bio) were plated at a density of 23 million cells per T-225 flask which was pre-coated with poly-l-lysine (Sigma-Aldrich). The cells were cultured in OptiMEM (Gibco) with 5% fetal bovine serum, 1% penicillin-streptomycin, 1% sodium pyruvate (Gibco), 1% MEM Non-Essential Amino Acids Solution (Gibco) and 1% HEPES (Gibco). Transfection was carried out using transfer plasmids along with second-generation lentiviral packaging plasmids, pMD2.G (Addgene) and psPAX2 (Addgene) using the Lipofectamine 3000 transfection reagent (Invitrogen). Six hours post transfection, the medium was replaced with OptiMEM (Gibco) containing 5% fetal bovine serum, 1% sodium pyruvate, 1% MEM Non-Essential Amino Acids Solution and 1% HEPES plus a viral boost reagent (Alstem) which was added according to manufacturer’s instructions. Viral supernatants were harvested 24 and 48 hours post transfection and centrifuged at 300g for 10 minutes at 4^*°*^C to remove any cell debris. Lentiviral particles were then concentrated using Lenti-X concentrator (Takara Bio) and stored overnight at 4^*°*^C. Next day, the virus was centrifuged at 1500g for 45 min at 4^*°*^C, resuspended in PBS at a concentration of 100X the original volume, and stored at −80^*°*^C. For T cell transduction, concentrated lentivirus was added directly to the T cells 24 hours after activation with Dynabeads Human T-Activator CD3/CD28 using 40 *µ*L virus per 1 ×10^6^ T cells in X-Vivo-15.

#### Generation of CRISPR knockout in primary human T cells by electroporation of Cas9–RNP

48 hours post of activation, electroporation was conducted using the Amaxa P3 Primary Cell 96-well 4D-Nucleofector Kit (Lonza). Lyophilized crRNA and tracrRNA from Dharmacon were reconstituted in nuclease-free duplex buffer (IDT) at a concentration of 160 *µ*M. Control T cells were targeted using the AAVS1 sequence GGGCCACTAGGGACAGGAT, RASA2-edited T cells with the sequence AGATATCACACATTACAGTG, and CUL5-edited T cells with the sequence ATTGGAGTAAGAGAATCCTA. crRNAs and tracrRNAs were then mixed 1:1 by volume and incubated for 30 minutes at 37C to form sgRNAs. Cas9 (stock concentration of 40 *µ*M, QB3 Macrolab) was then mixed with the sgRNAs 1:1 by volume for 15 minutes at 37^*°*^C to produce ribonucleoproteins (RNPs). T cells were resuspended in P3 buffer (Lonza) at a density of 1 ×10^6^ per 20 *µ*l, and subsequently combined with 3 *µ*l RNPs before being transferred to a 96-well electroporation plate (Lonza). Electroporation was performed using the pulse code EH115 on a 4D-Nucleofector 96-well Unit (Lonza) and cells were rescued by adding 80 *µ*l of X-VIVO-15 plus supplements for 15 minutes at 37C. The cells were then transferred to culture vessels containing X-Vivo-15 medium supplemented with IL-2 containing 100 IU per ml.

#### Determination of TCR+ T cells via flow cytometry

T cells were centrifuged for 5min at 300g and washed with 200 *µ*l of cell staining buffer consisting of PBS (Gibco) and 2% fetal bovine serum. The cells were then stained with Dextramer-HLAA*0201/SLLMWITQV-APC (Immudex) using 5 *µ*l of antibody in 50 *µ*l staining buffer for 10 min at room temperature in the dark. T cells were then washed twice in staining buffer and read on the Attune NXT Cytometer (Invitrogen). The data were analyzed with the software FlowJo version 10.9.

#### Repetitive stimulation assay

Tumor cells were maintained in complete RPMI, consisting of RPMI (Gibco), 1% penicillin-streptomycin (Gibco), GlutaMAX supplement (Gibco) and 10% fetal bovine serum (Corning). For the repetitive stimulation assay, tumor cells were split and plated in X-VIVO-15 plus supplements and 100 IU/mL of IL-2 in appropriate culturing vessels. T cells were then counted with a Cellaca MX High-throughput Cell Counter (Revvity), percentage of TCR+ cells was determined via flow cytometry and T cells were added onto fresh tumor cells every 48 hours maintaining a 1:1 effector-to-target ratio.

#### In vitro cytotoxicity assay using TCR T cells

Antigen-specific T cells were co-cultured in X-VIVO-15 plus supplements, 100 IU IL-2 per ml and 1X Glucose (Gibco) with mKate+ A375 cells that were pre-seeded in a 96-well flat-bottom plates. T cells were added in various E:T ratios. Over a 72-hour span, images were captured every 4 minutes using the Incucyte S3 live-cell imaging platform (Essen Bioscience), and the mKate+ object counts for each well were recorded over time.

### Occident pipeline application

#### Cell segmentation and tracking

Analysis was restricted to the 600 ×600 pixel center region of frames 50 to 350 (200 to 1400 minutes post co-culture). The small region ensured models could capture the majority of cells. The first 50 frames were skipped due to microscope lens artifacts. After 350 frames the density of cells becomes too high for separating individual cells. Each frame consists of brightfield and red nuclear fluorescent channel imaged at 10X resolution, at a scale of 1.245*µ*m/pixel. Each frame was first segmented using DeepCell nuclear segmentation model and cytoplasm segmentation model with micron per pixel set to 1.5 and 7.5 respectively^73^. DeepCell cytoplasm segmentation was post processed to fill any holes in segmentation and remove artifacts and debris segmented with less than 10 pixels. DeepCell nuclei segmentation was post-processed to remove nuclei segmentation fluorescent artifacts less than 80 pixels.

DeepCell fluorescent nuclei segmentation is near-perfect, so we further leveraged the information on two channels to refine cancer cells segmentation. By prompting nuclei centroids into Segment Anything Model (SAM) with default SAMPredictor parameters, we efficiently refined the boundaries of cancer blobs, as well as reducing the false positively detected T cell clumps that are treated as cancer cells due to the size threshold but actually lack a nuclei fluorescent signal. With cancer cell masks generated, T cell masks were subsequently rendered by subtracting cancer cell masks from DeepCell cytoplasm segmentation on the brightfield channel followed by thresholding T cells as objects less than 200 pixels.

As some of the T cells are missing from the DeepCell segmentation, we further leveraged Segment Anything Model (SAM), which is good at the general segmentation tasks, to help capture the missed T cells. SAM with ViT-B default weights was applied to segment cytoplasm images and T cells in between 10 to 200 pixels are added to the T cell masks.

With T cell masks and cancer cell masks generated across the time-lapse, these cells were tracked across frames using an in-house actualization of the tracking algorithm^74^ that ensures consistent identities. To tracking and detect division events of cancer nuclei, Caliban is used as part of the DeepCell cell tracking function with default settings.

#### Phenotype calculation

Segmentation returned five sets of cell masks - all T cells, all cancer cells, detached T cells, cancer cell aggregates, and cancer cell nuclei. For a set of cell masks, each frame consisted of a 600 ×600 array filled with either a zero, indicating no cell detected, or the unique cell ID number that occupies that region of the image.

#### Count and growth rates

The number of T cells and cancer cells at any time, *n*_*t*_, was calculated by counting the number of unique non-zero cell IDs in any frame. The T cell growth rate was calculated by fitting an exponential model log *n*_*t*_ = *a*_0_ + *a*_1_ ×*t*, where *t* is the time in minutes post co-culture, for each genetic perturbation across all wells. Statistical significance of difference between perturbations was checked by calculating the p-values of the interaction term for each pair of perturbations, *a*_*i*_, in a joint fit log *n*_*t*_ = *a*_0_ + *a*_1_×*t* + *a*_2_ ×*p*_1_ + *a*_*i*_×*t* ×*p*_1_, where *p*_1_ is a binary indicator if the sample comes from the control or test group. Cancer cell death rate was calculated by fitting a linear model *n*_*t*_ = *a*_*o*_ + *a*_1_×*t*, and statistical significance of differences between perturbation was calculated from the p-values of interaction terms in the joint fit *n*_*t*_ = *a*_0_ + *a*_1_×*t* + *a*_2_ ×*p*_1_ + *a*_*i*_×*t*×*p*_1_ as before.

#### Area, roundness, and velocity

Area for any cell type was calculated by counting the number of pixels in an individual cell’s mask at a given time. Perimeter was calculated by counting the number of zero or different cell ID entries adjacent (1 row above or below, 1 column left or right, no diagonals) to the mask. Roundness was determined by the ratio 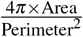. This ratio is 1 for a perfect circle and goes down as far as 0 for highly non-circular shapes. Speed was calculated as the euclidean distance between the centroid of a cell’s mask at times *t* and *t* + 1. Speed was not calculated for instances where the cell mask moves out of frame or disappears.

#### Cell divisions

Cancer division events were identified by Caliban from nuclei segmentation. The per T cell effect was calculated using a logistic regression fit 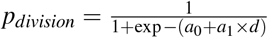, where *d* is the average local T cell density across the cancer cell’s track. The local T cell density for a cancer cell *c* is calculated by the average kernel distance to all other T cells detected in the frame at time *t*, 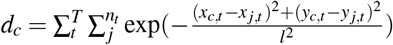, where (*x*_*i,t*_, *y*_*i,t*_) represents the centroid of cell *i* at time *t* in pixels. The length scale *l* was set to 30 pixels to match the approximate radius of a single cancer cell.

#### Interaction detection

An interaction was defined as having at least one pixel of the T cell mask adjacent to or overlapped with the cancer cell mask. Analysis was performed on interactions where both the T cell and cancer cell involved were tracked for at least two frames post-interaction. Area, roundness, and velocity were calculated as before. The interaction length was defined as the number of frames where both T cell and cancer cell involved met the interaction criteria.

#### Markov transition matrix estimation

Each frame of segmentation was split into 50 pixels ×50 pixels adjacent, non-overlapping regions (12 ×12 or 144 boxes total). For each time *t*, the state *s* of region *r* was determined by counting the number of individual T cells, *τ* and cancer cell, *κ*, centroids in the region. The Markov transition matrix for any starting state, *s*_0_, for each perturbation was estimated by summing the number of times any state *s*_*i*_ occurred in the same region *r* at *t* + 1 when the region *r* was at state *s*_0_ previously. This occurrence matrix is normalized by the total number of events to get a probability density. The probability of detachment, *p*_*d*_, was calculated as the total probability of any state with *τ* = 0 T cells and any number of cancer cells *κ* after an original state of (*τ* = 1, *κ* = 1). The time to detachment *l* was estimated by mean of a negative binomial distribution requiring *r* = 1 one success (one detachment), 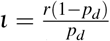, for each perturbation condition. Given all interactions were at least two frames in previous analysis, it was assumed that eight minutes of interaction had already that was added to the detachment time *l* to get the final interaction time. The recruitment probability was calculated by counting the number of times the unique id of the additional T cell in state *s*_*t*+1_ = (*τ* = 2, *κ* = 1) from *s*_*t*_ = (*τ* = 1, *κ* = 1) was also identified at *t*, divided by the total number of occurrence of *s*_*t*_. The proliferation probability was calculated as the difference between overall transition probability from (*τ* = 1, *κ* = 1) to (*τ* = 2, *κ* = 1) and the recruitment probability, equivalently the percentage of times the additional cell did not have a track at the previous time point.

## Author contributions statement

BEE, AM, SJ, CW, CY, and AV conceived Occident. MS, EM, WG, and DVV conceived Caliban.

MS, EM, CY, and GM developed the HITL labeling methodology. DVV, WG, EB, EM, and MS developed the deep learning cell-tracking methodology. MS, EM, EB, RD developed the cell-tracking and benchmarking software. MS and DVV developed the deep-learning nuclear segmentation methodology. CY developed whole-cell segmentation methodology for Occident. GM, TD, EB, and WG developed DeepCell Label and adapted it to live-cell images. GM developed new labeling tools within DeepCell Label to accelerate labeling. MS and WG developed the data-versioning methodology with DVC. MS, EM, GM, EP, and several unnamed image labelers developed the DynamicNuclearNet. CY and several unnamed image labelers created whole-cell labels for Occident. CY developed the image analysis pipelines used to analyze the data in this work. WG and DVV oversaw software engineering for DeepCell Label and Caliban. MS and DVV wrote the Caliban sections of the manuscript, with input from all authors.

SB, CC, and NK performed T cell experiments and collected live cell imaging data. CY performed cell segmentation and tracking with DeepCell, SAM, and Caliban. AV and IL implemented and executed Occident pipeline and analysis. AV and BEE analyzed the results from Occident. AV wrote the manuscript. DVV, AM, and JC and BEE supervised the project.

## Acknowledgments

The authors would like to acknowledge the incredible work of Tami Tolpa in creating the figures in this manuscript. We would also like to acknowledge Eric Ham, Leo Epstein, and Mihir Borkar for help with initial data exploration.

## Competing Interest Declaration

A.M. is a cofounder of Site Tx, Arsenal Biosciences, Spotlight Therapeutics and Survey Genomics, serves on the boards of directors at Site Tx, Spotlight Therapeutics and Survey Genomics, is a member of the scientific advisory boards of Site Tx, Arsenal Biosciences, Cellanome, Spotlight Therapeutics, Survey Genomics, NewLimit, Amgen, and Tenaya, owns stock in Arsenal Biosciences, Site Tx, Cellanome, Spotlight Therapeutics, NewLimit, Survey Genomics, Tenaya and Lightcast and has received fees from Site Tx, Arsenal Biosciences, Cellanome, Spotlight Therapeutics, NewLimit, Gilead, Pfizer, 23andMe, PACT Pharma, Juno Therapeutics, Tenaya, Lightcast, Trizell, Vertex, Merck, Amgen, Genentech, GLG, ClearView Healthcare, AlphaSights, Rupert Case Management, Bernstein and ALDA. A.M. is an investor in and informal advisor to Offline Ventures and a client of EPIQ. The Marson laboratory has received research support from the Parker Institute for Cancer Immunotherapy, the Emerson Collective, Arc Institute, Juno Therapeutics, Epinomics, Sanofi, GlaxoSmithKline, Gilead and Anthem and reagents from Genscript and Illumina. D.V.V. is the scientific founder of Aizen Therapeutics and holds equity in the company. B.E.E. is on the Scientific Advisory Board for ArrePath Inc, Crayon Bio, and Freenome; she consults for Neumora. B.E.E. and A.V. were funded in part by Helmsley Trust grant AWD1006624, NIH NCI 5U2CCA233195, the Parker Institute for Cancer Immunotherapy (PICI), NHGRI R01 HG013736, and NIH NHGRI R01 HG012967. BEE is a CIFAR Fellow in the Multiscale Human Program. A.V. has received fees from OpenProtein.AI. All other authors declare no competing interests.

## Supplement

Supplementary File 1: DeepCell Label User Manual

Supplementary File 2: Segmentation Correction Instructions

Supplementary File 3: Tracking Correction Instructions

**Supplementary Table 1.** Publicly available labeled datasets for two-dimensional temporal (2D+T) cell tracking. CTMC: Cell Tracking with Mitosis Detection Dataset Challenge, CTC: Cell Tracking Challenge, Fluor: fluorescent, DIC: differential interference contrast, WC: whole cell, Part: partial.

| Name | Modality | Annotation Type | Cell Types | Objects | Tracks | Divisions | Source |
| --- | --- | --- | --- | --- | --- | --- | --- |
| DynamicNuclearNet | Fluor. Nuclei | Nuclear Mask | 5 | 647,322 | 16,501 | 2,621 | This work |
| DeepSea | Phase | Nuclear Mask | 3 | 100,000 | 2,576 | 137 | Zargari *et. al.*<sup>34</sup> |
| BTrack/CellX | Fluor. Nuclei | Nuclear Mask | 1 | - | 1,032 | - | Cuny *et. al.*<sup>72</sup> |
| CTMC | DIC | Bounding Box | 14 | 2,045,834 | 2,900 | 457 | Anjum & Gurrari<sup>30</sup> |
| C2C12 | Phase | Centroid | 1 | 135,859 | 23,400 | 7,159 | Kerr *et. al.*<sup>28</sup> |
| CTC 2D+T | DIC | Part. WC Mask | 1 | - | 70 | 18 | CTC<sup>75</sup> |
| CTC 2D+T | Phase | Part. WC Mask | 2 | - | 2,415 | 1,019 | CTC<sup>75</sup> |
| CTC 2D+T | Fluor. Nuclei | Part. Nuclear Mask | 3 | - | 1,227 | 395 | CTC<sup>75</sup> |
| CTC 2D+T | Fluor. WC | Part. WC Mask | 2 | - | 128 | 10 | CTC<sup>75</sup> |
| CTC 2D+T | Brightfield | Part. WC Mask | 2 | - | 459 | 242 | CTC<sup>75</sup> |

**Supplementary Table 2.** Benchmarking the performance of different tracking methods on the test split of DynamicNuclearNet. Bold font indicates the best scores on predicted segmentations. Italic font denotes the best scores on ground truth (GT) segmentations. CTC: Cell Tracking Challenge, DET: CTC detection accuracy^46^, SEG: CTC segmentation accuracy^76^, TRA: CTC tracking accuracy^76^, Div R: division recall, Div P: division precision, Div F1: division F1 score, MBC: mitotic branching correctness^33^, AA: association accuracy^47,48^, TE: target efficiency^47,48^.

| Tracking | Segmentation | DET | SEG | TRA | Div R | Div P | Div F1 | MBC | AA | TE |
| --- | --- | --- | --- | --- | --- | --- | --- | --- | --- | --- |
| Caliban | Caliban | **0.990** | **0.913** | **0.989** | **0.90** | **0.77** | **0.83** | **0.71** | 0.95 | 0.96 |
|  | GT | *1.000* | *1.000* | *0.999* | *0.95* | *0.92* | *0.94* | *0.88* | 0.98 | 0.98 |
| Baxter | Caliban | 0.988 | 0.908 | 0.986 | 0.49 | 0.73 | 0.59 | 0.41 | **0.98** | **0.98** |
|  | GT | 0.997 | 0.996 | 0.997 | 0.60 | 0.89 | 0.72 | 0.56 | *1.00* | *0.99* |
| Trac<sup>x</sup> | Caliban | **0.990** | **0.913** | **0.989** | 0.35 | 0.29 | 0.32 | 0.19 | 0.95 | 0.95 |
|  | GT | *1.000* | *1.000* | *0.999* | 0.34 | 0.54 | 0.42 | 0.27 | 0.98 | 0.98 |
| CellTrackerGNN | CellTrackerGNN | 0.815 | 0.682 | 0.812 | 0.43 | 0.05 | 0.08 | 0.04 | 0.76 | 0.76 |
|  | GT | *1.000* | 0.999 | 0.996 | 0.65 | 0.10 | 0.18 | 0.10 | 0.93 | 0.93 |
| EmbedTrack | EmbedTrack | 0.816 | 0.642 | 0.815 | 0.64 | 0.17 | 0.27 | 0.15 | 0.82 | 0.82 |

**Supplementary Table 3.**
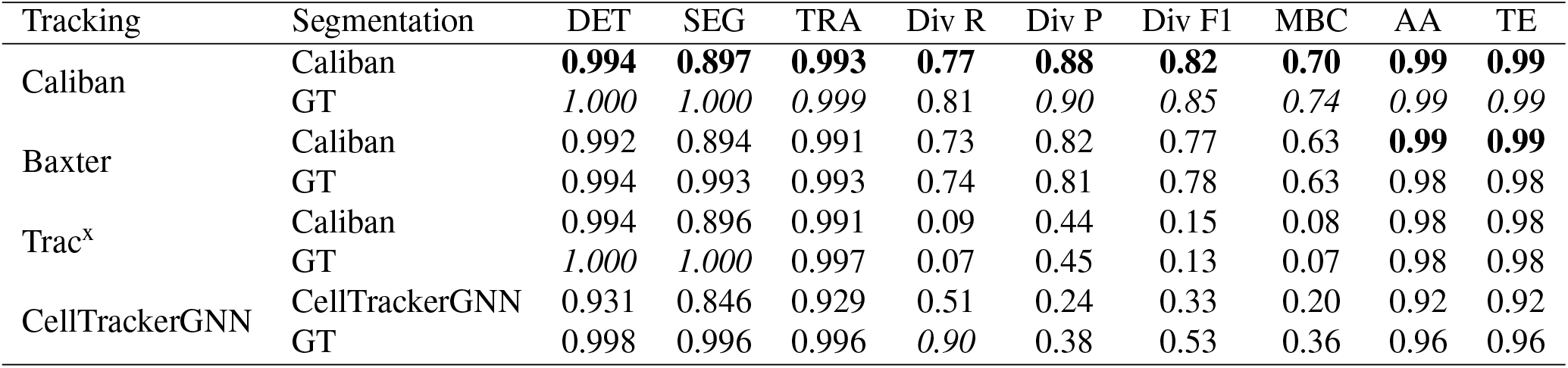
Benchmarking the performance of different tracking methods on the CTC Fluo-N2DL-HeLa test movies. Bold font indicates the best scores on predicted segmentations. Italic font denotes the best scores on ground truth (GT) segmentations. CTC: Cell Tracking Challenge, DET: CTC detection accuracy^46^, SEG: CTC segmentation accuracy^76^, TRA: CTC tracking accuracy^76^, Div R: division recall, Div P: division precision, Div F1: division F1 score, MBC: mitotic branching correctness^33^, AA: association accuracy^47,48^, TE: target efficiency^47,48^.

| Tracking | Segmentation | DET | SEG | TRA | Div R | Div P | Div F1 | MBC | AA | TE |
| --- | --- | --- | --- | --- | --- | --- | --- | --- | --- | --- |
| Caliban | Caliban | **0.994** | **0.897** | **0.993** | **0.77** | **0.88** | **0.82** | **0.70** | **0.99** | **0.99** |
|  | GT | *1.000* | *1.000* | *0.999* | 0.81 | *0.90* | *0.85* | *0.74* | *0.99* | *0.99* |
| Baxter | Caliban | 0.992 | 0.894 | 0.991 | 0.73 | 0.82 | 0.77 | 0.63 | **0.99** | **0.99** |
|  | GT | 0.994 | 0.993 | 0.993 | 0.74 | 0.81 | 0.78 | 0.63 | 0.98 | 0.98 |
| Trac<sup>x</sup> | Caliban | 0.994 | 0.896 | 0.991 | 0.09 | 0.44 | 0.15 | 0.08 | 0.98 | 0.98 |
|  | GT | *1.000* | *1.000* | *0.997* | 0.07 | 0.45 | 0.13 | 0.07 | 0.98 | 0.98 |
| CellTrackerGNN | CellTrackerGNN | 0.931 | 0.846 | 0.929 | 0.51 | 0.24 | 0.33 | 0.20 | 0.92 | 0.92 |
|  | GT | 0.998 | 0.996 | 0.996 | *0.90* | 0.38 | 0.53 | 0.36 | 0.96 | 0.96 |

**Supplementary Figure 1.**
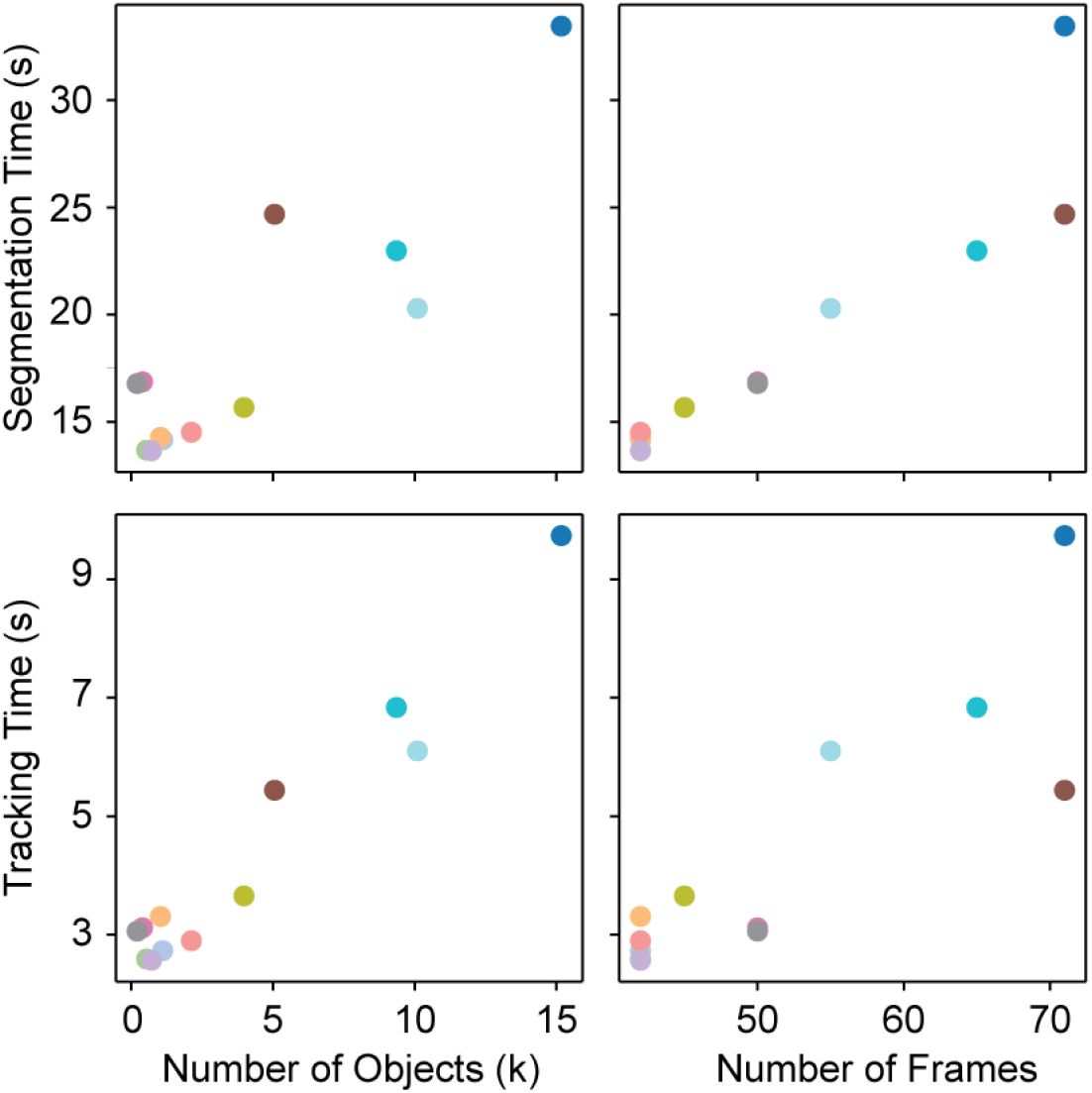
Runtime for segmentation and tracking with Caliban. The total runtime for segmentation and tracking is plotted as a function of the number of objects and frames in the sample. Each point represents a movie in the test data split, with a unique color assigned to each movie.

**Supplementary Figure 2.**
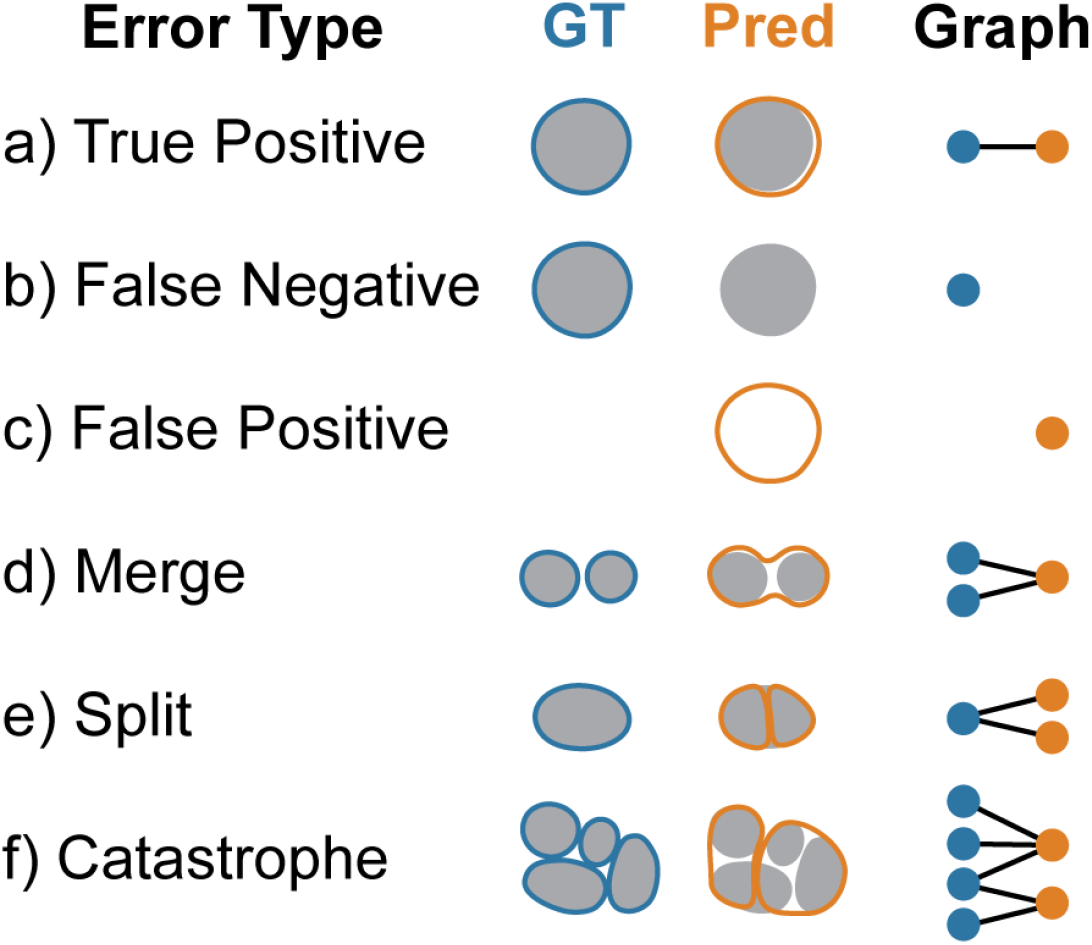
Object-based evaluation of segmentation performance. Segmentation predictions were evaluated based on object-level accuracy by first constructing a graph in which edges indicate an overlap between two objects. Each subgraph is then isolated and analyzed to identify the type of segmentation error present. (a) Subgraphs with one ground truth (GT) and one predicted node represent a true positive segmentation. Subgraphs containing only one node represent (b) a false negative if the node is ground truth or (c) a false positive if the node is predicted. Subgraphs with three nodes indicate (d) a merge if two ground truth nodes are associated with one predicted node or (e) a split if two predicted nodes are associated with one ground truth node. (f) Finally, all subgraphs containing more than three nodes are assigned to the catastrophe error class.

**Supplementary Figure 3.**
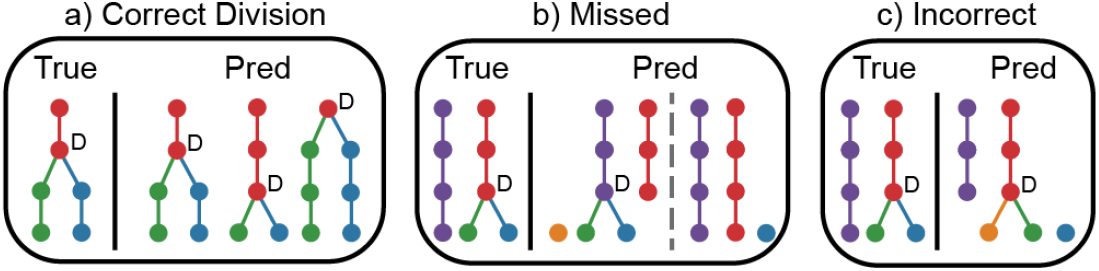
Division-based evaluation of tracking performance. Division events are classified as correct, missed, or incorrect based on a comparison of the true and predicted tracking graphs. (a) A division is considered correct if the prediction links the parent to the correct daughters within one frame of the ground truth division event. We allow divisions to shift in time because segmentation predictions can change when the cell is identified as one or two objects. (b) Divisions are identified as missed if the daughter cells are assigned to the incorrect parent or if no parent is identified. (c) A division is incorrect if the parent is assigned to only one of the correct daughter cells.

**Supplementary Figure 4.**
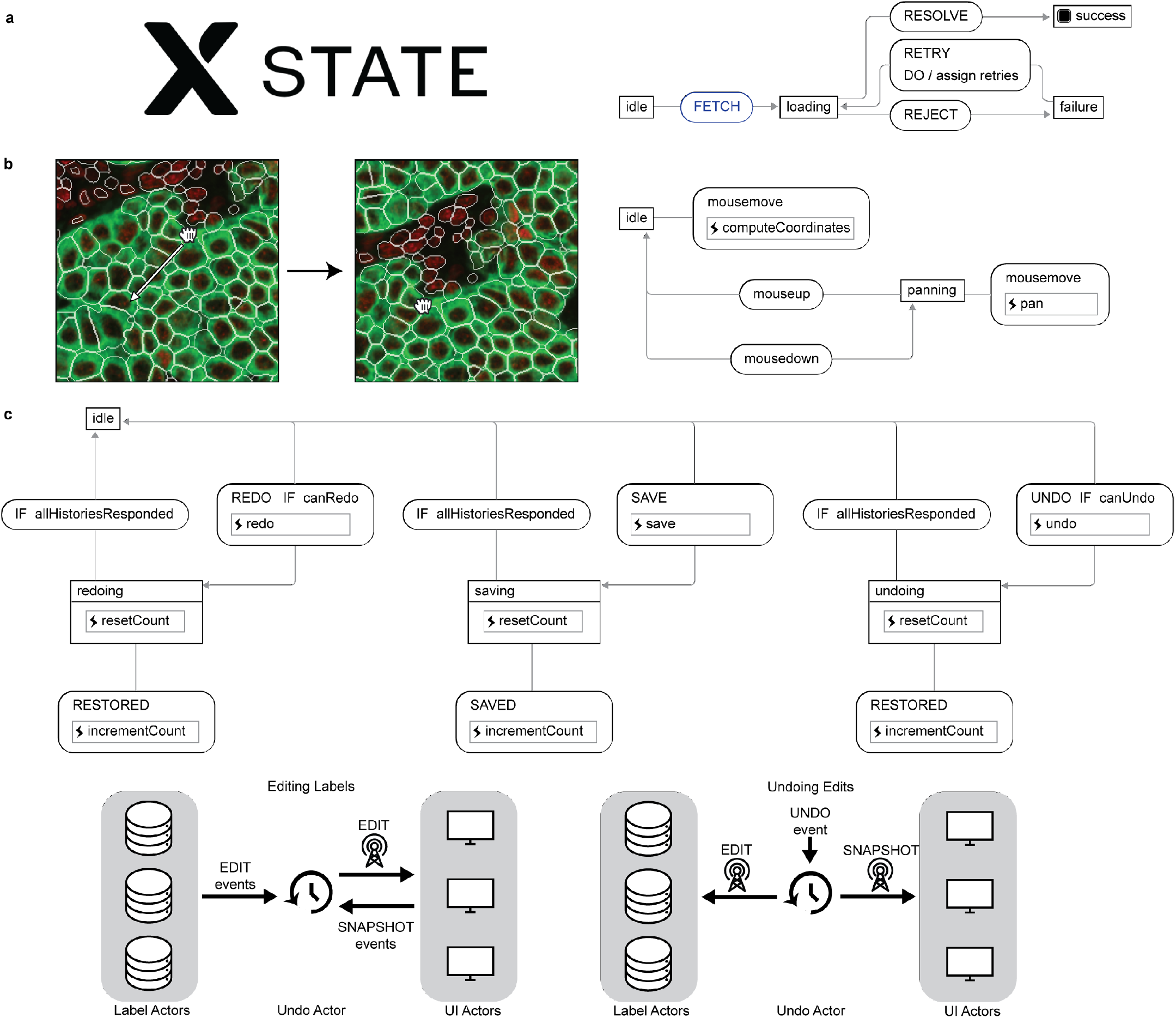
State management in the DeepCell Label backend. (a) DeepCell Label uses XState, a library for event-driven programming and state management, to drive its user interface (UI) and data labeling logic. Actors defined with XState control interactions with UI components and with labeled data. Actors consist of a context with arbitrary data, such as the settings for UI component or data labels; and a set of finite states and transitions between them. Actors receive events, which trigger transitions between states. Events are sent by user interactions or sent from other actors. (b) The panning states for the canvas actor. The actor begins in an idle state, where mousemove events update the position of the cursor. Upon a mousedown event, the actor transitions into a panning state, where mousemove events instead change the position of the image, enabling the user to browse the image. Once the user releases the mouse and triggers a mouseup event, the actor returns to the idle state. (c) The undo actor broadly orchestrates state across all UI and data labeling actors to enable undoing and redoing edits to labeled data. Actors that wish to subscribe to undo and redo events send an event to the undo actor to register itself as a UI actor or a data labeling actor. When data labeling actors edit their labels, they send a SNAPSHOT event to the undo actor, which then collects snapshots of all registered UI actors. When the user sends an undo event to the undo actor, the undo actor resends the snapshot events associated with the edit, globally restoring the application to its state just before the edit. Each registered actor maintains its own state and logic on how to implement undoable behavior, while the undo actor serves as a shared channel to coordinate registered actors.

**Supplementary Figure 5.**
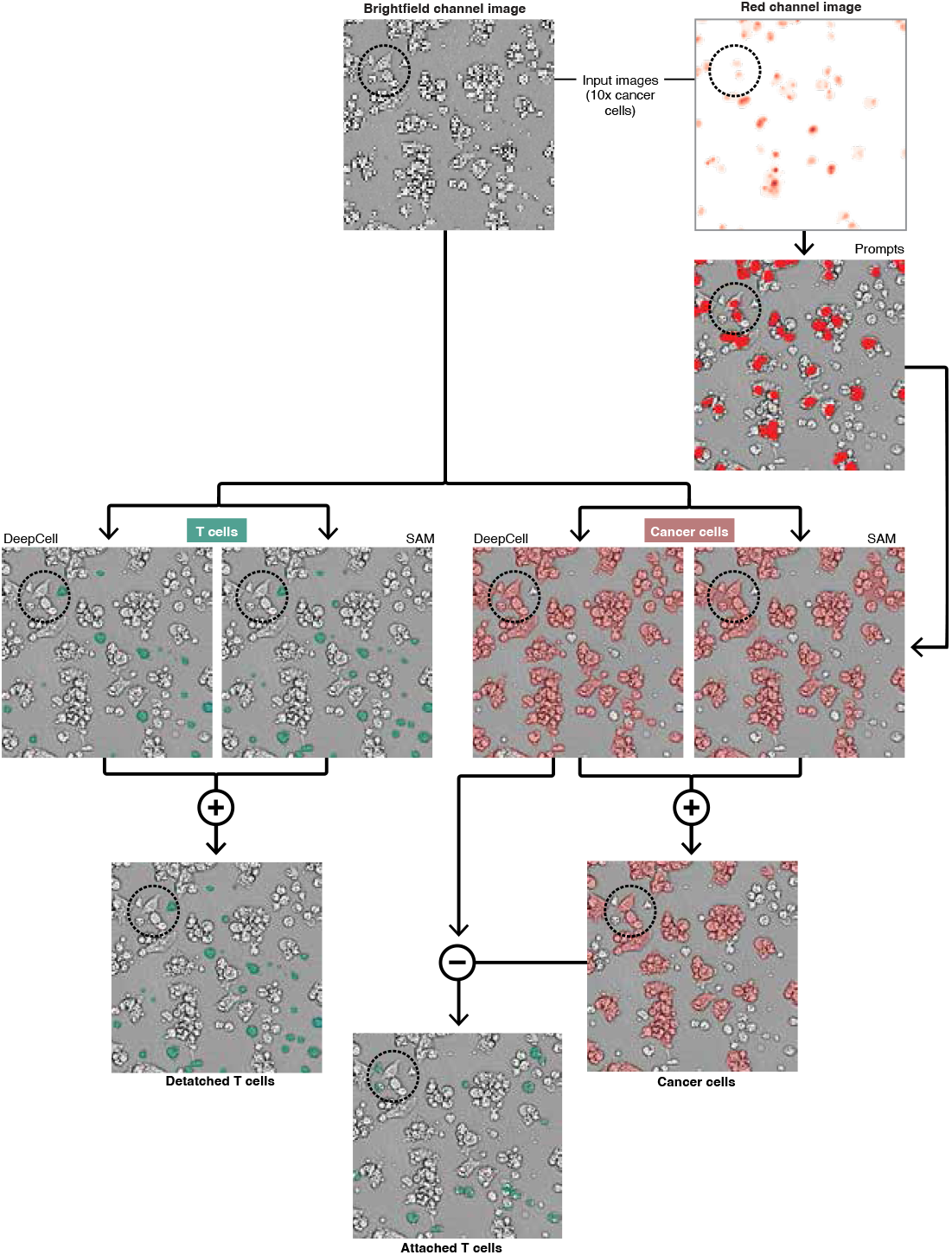
Segmentation pipeline for special cell type masks. A combination of Caliban and SAM was used to get special cancer cell and T cell masks. Red nuclear channel images were used to generate cancer nuclei masks. Phase images were used to generate T cell masks with DeepCell/Caliban and SAM, then ensembled for final masks. Red and phase images were used to generate cancer cell aggregate masks with DeepCell/Caliban. Nuclear masks were used to prompt SAM to generate a second set of masks ensembled with DeepCell to generate individual cancer cell masks. The difference between the aggregate masks and individual cancer masks generated additional attached T cells for inclusion with the previous T cell masks.

## Notes

https://github.com/vanvalenlab/Caliban-2024_Schwartz_et_al

https://github.com/bee-hive/Occident-Paper/tree/main

